# Systemic Autoimmunity Remodels Microbiota-Reactive CD4⁺ T Cell Immunity

**DOI:** 10.64898/2026.10.17.759871

**Authors:** Maricielo J. Vaella-Alarcon, Miki Kume, Silvia R. Davila-Paico, Luis F. Soto-Ugaldi, Katherine S. Saldana-Morales, Gillian L. McClennen, Jin Din, Samantha St Jean, Daniel F. Zegarra-Ruiz

**Author notes:** Corresponding author. (D.F.Z.-R.). These authors contributed equally to this work (Maricielo J. Vaella-Alarcon, Miki Kume).

## Abstract

The gut microbiota contributes to autoimmune disease, yet how systemic autoimmunity reshapes microbiota-reactive CD4⁺ T cell responses and how these cells influence disease progression remain poorly understood. Previous studies have linked microbiota-reactive T cells and lupus-associated pathobionts to disease pathogenesis, but the pathways that regulate the accumulation, differentiation, and pathogenicity of microbiota-reactive T cells during autoimmunity remain unclear. Here, we show that systemic autoimmunity drives the expansion, accumulation, and functional differentiation of microbiota-reactive CD4⁺ T cells. Using segmented filamentous bacteria tetramers, we identify antigen-defined microbiota-reactive CD4⁺ T cells that undergo systemic accumulation and Th1/Tfh polarization during disease. We further identify reactivation of intestinal dendritic cell trafficking as a mechanism linking systemic autoimmunity to the accumulation of microbiota-reactive CD4⁺ T cells in peripheral tissues. In contrast to commensal-reactive T cells, CD4⁺ T cells conditioned by the lupus-associated pathobiont *Enterococcus gallinarum* markedly exacerbate autoimmunity following adoptive transfer through IL-21- and IFNγ-dependent mechanisms. Together, these findings demonstrate that systemic autoimmunity reshapes microbiota-reactive CD4⁺ T cell immunity and establish microbial context as a critical determinant of microbiota-reactive CD4⁺ T cell pathogenicity.

**SIGNIFICANCE STATEMENT:** Microbiota-reactive CD4⁺ T cells contribute to host defense and have been implicated in autoimmune diseases, but how systemic autoimmunity alters these responses has remained unclear. We show that systemic autoimmunity promotes the expansion, accumulation, and differentiation of microbiota-reactive CD4⁺ T cells by reactivating intestinal dendritic cell trafficking. We further demonstrate that microbial context determines the pathogenic potential of these cells, with pathobiont-conditioned CD4⁺ T cells acquiring disease-promoting activity. These findings identify a mechanism by which chronic autoimmune inflammation reshapes host-microbiota adaptive immunity and influences disease progression.

## INTRODUCTION

The gut microbiota plays a fundamental role in shaping immune responses, and disruption of host-microbiota interactions has been linked to autoimmune diseases such as multiple sclerosis, type 1 diabetes, and systemic lupus erythematosus (SLE), contributing to chronic inflammation and immune dysfunction (*1–3*). Human and mouse studies have identified specific intestinal microbes that become enriched during autoimmune disease and contribute to disease progression through distinct mechanisms (*2, 4–8*). Pathobionts such as *Enterococcus gallinarum* and *Lactobacillus reuteri* induce inflammatory CD4⁺ T cell responses and exacerbate disease, whereas commensal microbes expressing orthologs of the lupus autoantigen Ro60 activate microbiota-reactive T cells that cross-react with self-antigens (*4, 5, 9, 10*). Together, these findings highlight microbiota-reactive CD4⁺ T cells as potential mediators of autoimmune disease progression. However, the immune mechanisms through which systemic autoimmunity reshapes microbiota-reactive CD4⁺ T cell responses remain poorly understood.

Microbiota-reactive CD4⁺ T cells are abundant components of the normal immune repertoire and are present in both circulation and peripheral tissues, where they contribute to host defense and immune homeostasis (*11*). Studies in humans have shown that these cells undergo functional alterations during chronic inflammatory disease, suggesting that inflammatory environments can reshape microbiota-reactive immunity (*12*). Despite growing evidence linking intestinal microbes to lupus through pathobiont expansion, microbial translocation, and molecular mimicry, it remains unclear how systemic autoimmunity reshapes microbiota-reactive CD4⁺ T cell responses (*4–6, 9*). Whether autoimmune disease promotes the accumulation and differentiation of microbiota-reactive T cells, whether these cells acquire pathogenic functions during disease, and why responses directed against distinct intestinal microbes result in different autoimmune outcomes remain fundamental unanswered questions.

Here, we used complementary models of TLR7-driven systemic autoimmunity to investigate how systemic autoimmunity reshapes microbiota-reactive CD4⁺ T cell immunity. We show that autoimmunity promotes the expansion, accumulation, and functional differentiation of these cells and identify reactivation of intestinal dendritic cell trafficking as a mechanism linking systemic autoimmunity to the accumulation of microbiota-reactive CD4⁺ T cells in peripheral tissues during disease. Finally, we demonstrate that microbial context dictates pathogenic potential, with pathobiont-conditioned, but not commensal-reactive, CD4⁺ T cells exacerbating autoimmune disease following adoptive transfer. Together, these findings identify mechanisms that regulate the accumulation, differentiation, and pathogenicity of microbiota-reactive CD4⁺ T cells during systemic autoimmunity.

## RESULTS

### TLR7-driven autoimmunity promotes microbiota-reactive CD4⁺ T cell responses

The gut microbiota contributes to autoimmune diseases, yet how systemic autoimmunity alters microbiota-reactive CD4⁺ T cell responses remains poorly understood. We hypothesized that TLR7-driven autoimmunity alters the magnitude and composition of microbiota-reactive CD4^+^ T cell responses. To assess microbiota-reactive T cell responses in an unbiased manner, we treated mice with the TLR7 agonist imiquimod (IMQ) for six weeks to induce systemic autoimmunity (*9, 13*) and stimulated mesenteric lymph node (MLN) and splenic cells ex vivo with autologous fecal lysates, using Nur77 induction as a readout of antigen-dependent TCR signaling (*14*) (Fig. 1A, Fig. S1A, B).

**Fig. 1.**
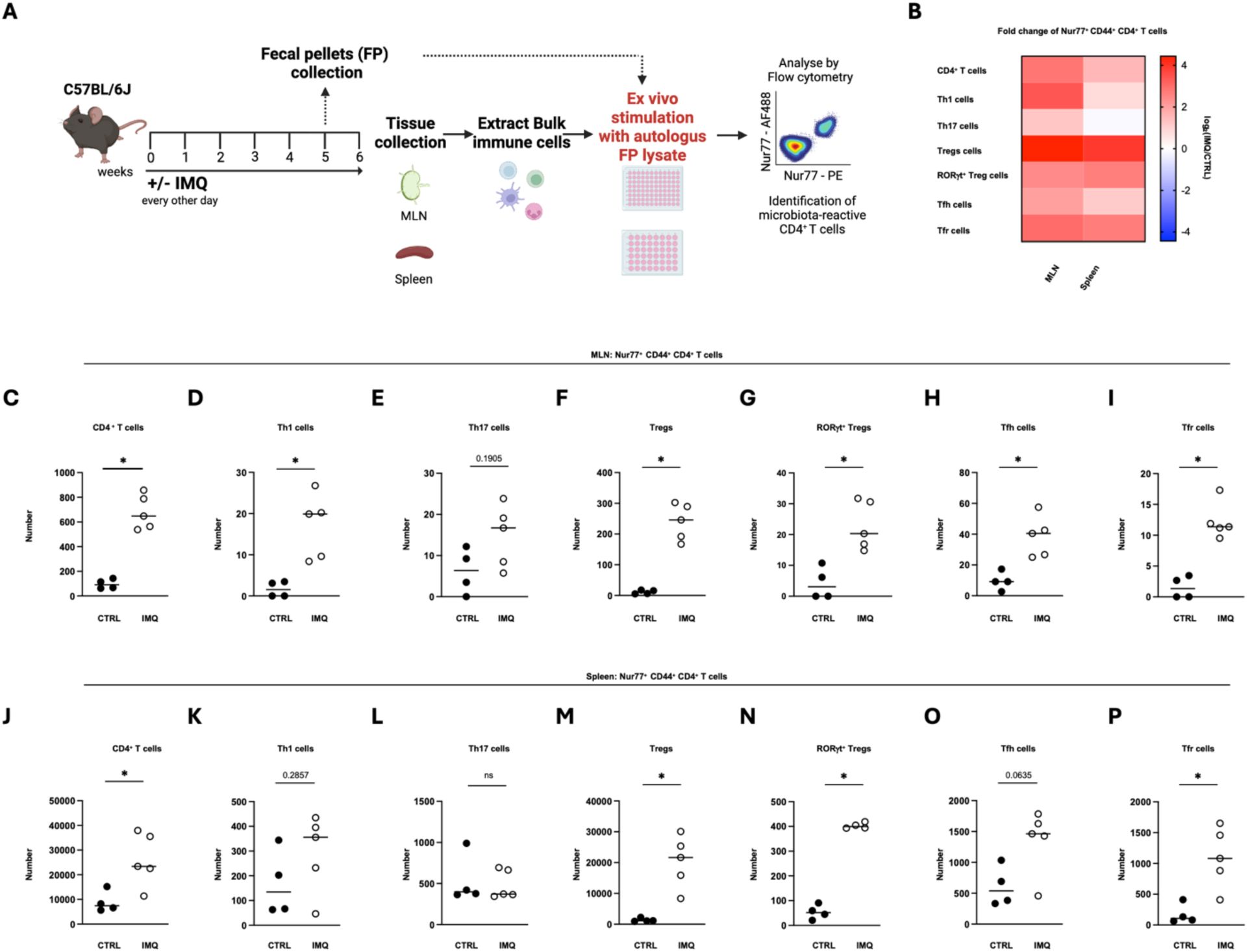
TLR7-driven autoimmunity promotes microbiota-reactive CD4⁺ T cell responses across lymphoid tissues. (A) Experimental schematic of ex vivo T cell restimulation with fecal bacterial lysate. (B) Heatmap depicts log_2_(IMQ/CTRL) fold change for Nur77⁺ CD44⁺ CD4⁺ T cells in the MLN and spleen of control (CTRL) and imiquimod (IMQ)-treated mice (n = 4-5). (C-P) Counts of Nur77-expressing CD44^+^ CD4^+^ (C, J), Th1 (D, K), Th17 (E, L), Treg (F, M), RORγt^+^Treg (G, N), Tfh (H, O), and Tfr (I, P) T cells in the mesenteric lymph nodes (C-I) or spleen (J-P) of CTRL and IMQ mice (n = 4-5). Panel A was created with Biorender.com. Data shown as means. Statistical significance was determined by unpaired two-tailed Mann-Whitney U test. *\*p < 0.05, **p < 0.01, ***p < 0.001, ****p < 0.0001.* See also Figure S1.

To define the impact of autoimmunity on microbiota-reactive CD4⁺ T cell responses, we quantified fecal lysate-induced Nur77 expression across CD4⁺ T cell subsets in IMQ-treated mice relative to controls. TLR7-driven autoimmunity promoted broad remodeling of microbiota-reactive CD44^+^ CD4⁺ T cell populations in both the MLN and spleen, with the greatest changes observed among regulatory CD4⁺ T cell subsets (Fig. 1B).

In the MLN, IMQ-treated mice exhibited an increased number of Nur77⁺CD44⁺ CD4⁺ T cells following fecal lysate stimulation, indicating the accumulation of microbiota-reactive T cells within gut-draining lymphoid tissues (Fig. 1C, Fig. S1C). This increase extended across multiple CD4⁺ T cell subsets, including Th1 cells, regulatory T cells (Tregs), RORγt^+^ Tregs, follicular helper T cells (Tfh), and follicular regulatory T cells (Tfr), demonstrating that autoimmunity broadly remodels the composition of microbiota-reactive CD4^+^ T cell populations rather than selectively expanding a single lineage (Fig. 1D-I, Fig. S1D-I).

A different pattern emerged in the spleen. IMQ-treated mice showed a reduced frequency but increased number of Nur77⁺CD44⁺ CD4⁺ T cells, indicating numerical accumulation of microbiota-reactive antigen-experienced CD4⁺ T cells despite changes in subset composition (Fig. 1J, Fig. S1J). Among these populations, microbiota-reactive Tregs increased in both frequency and number, whereas Tfh cells decreased in frequency but showed a strong trend toward increased numbers (Fig. 1K-P, Fig. S1K-P). Overall, CD44⁺ subset analyses revealed accumulation across multiple microbiota-reactive CD4⁺ T cell subsets despite more variable changes in frequency (Fig. S1K-P). Similar changes were observed when Nur77 responses were analyzed across total CD4⁺ T cells in both the MLN and spleen, although the magnitude of these responses was greater within the CD44⁺ antigen-experienced compartment (Fig. S1Q-AA). Together, these findings indicate that systemic autoimmunity promotes the accumulation of microbiota-reactive CD4⁺ T cells across multiple lineages, with the most pronounced changes occurring within antigen-experienced CD44⁺ populations.

### Antigen-defined microbiota-reactive CD4⁺ T cells accumulate and acquire effector phenotypes during autoimmunity

Because fecal lysate stimulation does not identify the microbial specificities responsible for these responses, we next used MHC class II tetramers loaded with an immunodominant peptide from the commensal gut microbe segmented filamentous bacteria (SFB) (I-A^b^/SFB3340) (*15, 16*) to determine whether TLR7-driven autoimmunity promotes the accumulation and differentiation of antigen-defined microbiota-reactive CD4⁺ T cells.

Adult SFB-free mice were treated with imiquimod for six weeks to induce systemic autoimmunity and colonized with SFB during the final two weeks of treatment (Fig. S2A). Consistent with previous reports (*9, 17*), IMQ-treated mice developed features of systemic autoimmunity, including splenomegaly, elevated autoantibodies, and type I interferon responses; however, introduction of SFB during the final two weeks of disease induction did not significantly alter these parameters within the timeframe examined (Fig. S2B-M). These findings establish SFB as a defined microbial antigen for tracking microbiota-reactive CD4⁺ T cells in this model while minimizing confounding effects on disease severity. Using I-A^b^/SFB3340 tetramers, we observed increased frequencies and numbers of SFB tetramer⁺ CD4⁺ T cells in the spleen of IMQ-treated mice compared with untreated controls (Fig. 2A, S3A). Hereafter, these cells are referred to as SFB tetramer⁺ T cells. Expansion of SFB tetramer⁺ T cells was also detected across multiple peripheral tissues, including MLN, Peyer’s patches, and small intestine, indicating systemic expansion and accumulation of antigen-defined microbiota-reactive CD4⁺ T cells during autoimmunity (Fig. 2B). In contrast to SFB tetramer⁺ T cells, CD4⁺ T cells specific for bacterial flagellin from an unrelated microorganism or the irrelevant control peptide CLIP showed no significant changes following IMQ treatment (Fig. S3B, C). Similar results were observed across different SFB colonization strategies, indicating that SFB tetramer⁺ T cell accumulation occurred regardless of whether colonization preceded or followed disease initiation (Fig. S3D, E). Consistent with active expansion, splenic SFB tetramer⁺ T cells exhibited increased Ki67 expression compared with tetramer-negative CD4⁺ T cells, indicating enhanced proliferation during autoimmunity (Fig. S3F) (*18*).

**Fig. 2.**
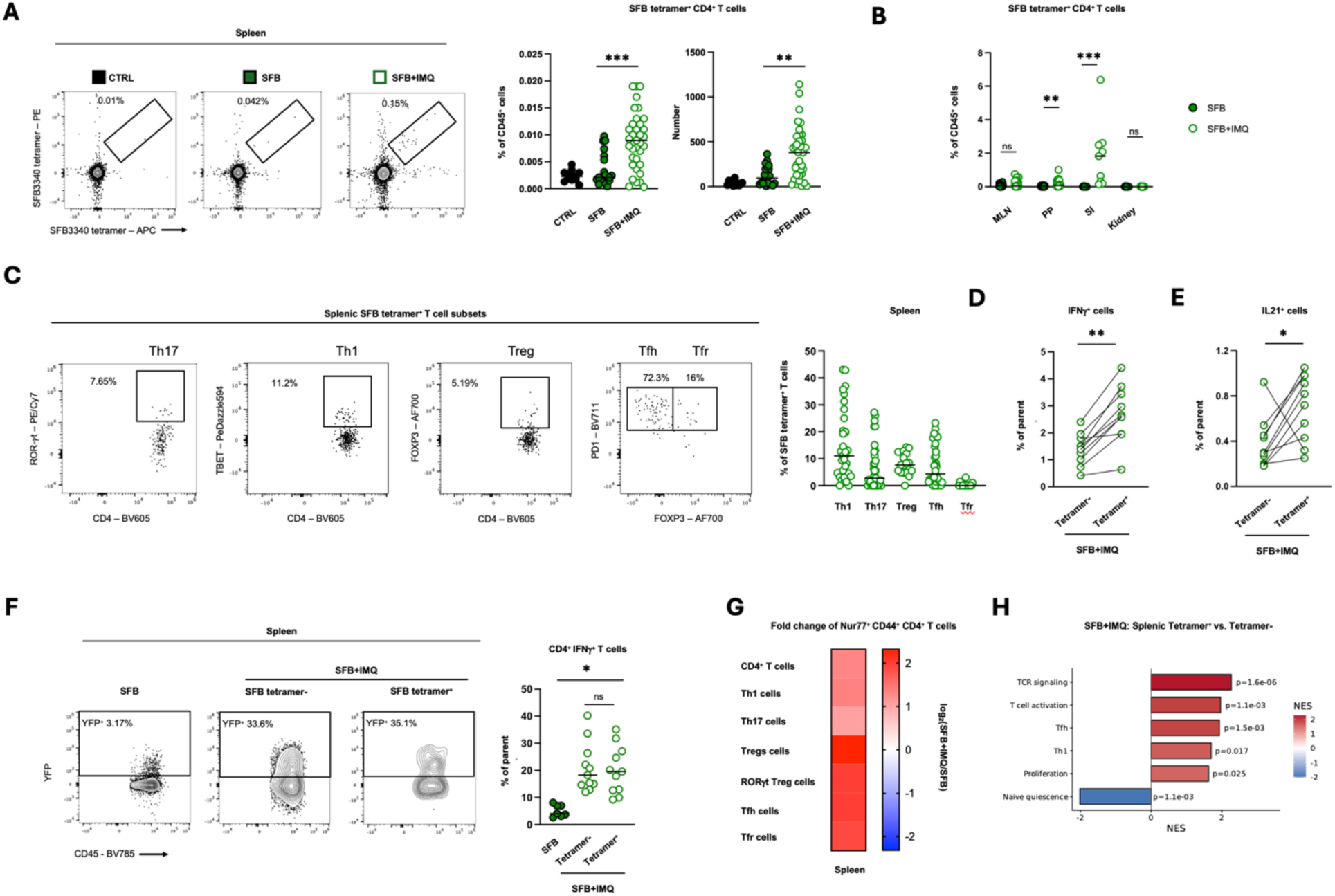
SFB tetramer⁺ CD4⁺ T cells expand and acquire Th1/Tfh programs during TLR7-driven autoimmunity. (A) SFB tetramer^+^ CD4⁺ T cells in the spleen of control (CTRL), SFB-colonized (SFB), or SFB-colonized imiquimod-treated (SFB+IMQ) mice (n = 12-40). (B) Frequency of SFB tetramer^+^ CD4⁺ T cells of total CD45^+^ cells in the mesenteric lymph node (MLN), Peyer’s patches (PP), small intestinal lamina propria (SI), and kidney of SFB and SFB+IMQ mice (n = 7-34). (C) Th1, Th17, Treg, Tfh, and Tfr subsets in splenic SFB tetramer^+^ CD4⁺ T cells (n = 17-35). (D-E) IFNγ (D) and IL-21 (E) production in SFB tetramer^-^ and SFB tetramer^+^ CD4⁺ T cells from spleen of SFB-colonized imiquimod-treated mice (n = 10). (F) In reporter mice expressing YFP at the IFNγ locus, frequency of IFNγ^+^ cells of total splenic CD4^+^ T cells from SFB, and SFB tetramer^-^ and SFB tetramer^+^ splenic CD4^+^ T cells from SFB+IMQ (n = 7-11). (G) Log_2_(SFB+IMQ/SFB) fold change of counts of Nur77⁺ CD44⁺ CD4⁺ T cell subsets in spleen of SFB and SFB+IMQ mice (n = 4-5). (H) Differentially expressed pathways between SFB tetramer^-^ and SFB tetramer^+^ CD4^+^ T cells from the spleen of SFB-colonized IMQ-treated mice. Data shown as means. Statistical significance was determined by Kruskal-Wallis test (A, F), unpaired two-tailed Mann-Whitney U test (B), or two-tailed Wilcoxon test (D-E). Statistical tests for RNA-seq are described in the sequencing analysis methods. *\*p < 0.05, **p < 0.01, ***p < 0.001, ****p < 0.0001.* See also Figures S2, S3, and S4.

To determine how autoimmunity influences the differentiation of antigen-defined microbiota-reactive CD4⁺ T cells, we examined SFB tetramer⁺ T cell subsets based on transcription factor expression across lymphoid and peripheral tissues. In the spleen, SFB tetramer⁺ T cells exhibited a predominantly Th1 phenotype, with smaller Treg and Tfh populations also detected, whereas Th17 cells represented a minor fraction of the antigen-reactive compartment (Fig. 2C). Analysis of SFB tetramer⁺ T cells across intestinal tissues revealed tissue-specific remodeling during autoimmunity. Relative to SFB-colonized control mice, SFB tetramer⁺ T cells in the MLN exhibited increased Th1 and Treg differentiation, whereas Peyer’s patches exhibited reduced Th17 frequencies and the small intestine remained largely unchanged (Fig. S3G). Analysis of total CD4⁺ T cells also revealed tissue-specific remodeling during autoimmunity. In the spleen, Th1 and Tfh populations were increased, while the MLN exhibited increased Th1 and Treg populations. In contrast, Peyer’s patches and the small intestine showed minimal changes, whereas the kidney displayed marked enrichment of Th1 and Th17 cells (Fig. S3H, I).

Consistent with the enrichment of Th1 and Tfh populations among SFB tetramer⁺ T cells, splenic SFB tetramer⁺ T cells produced elevated levels of IFNγ and IL-21 following ex vivo PMA/ionomycin stimulation compared with tetramer-negative T cells, further supporting acquisition of Th1 and Tfh effector programs during autoimmunity (Fig. 2D, E). Additional cytokine analyses revealed more limited changes in TNFα and IL-17A production, indicating that Th1 and Tfh programs represent the dominant effector states acquired by SFB tetramer⁺ T cells in this setting (Fig. S3J-L). Analysis of IFNγ reporter mice further supported in vivo Th1 polarization during TLR7-dependent autoimmunity, revealing increased IFNγ⁺ cells among both SFB tetramer⁺ and tetramer-negative CD4⁺ T cells (Fig. 2F).

We next asked whether SFB tetramer⁺ T cells remained responsive to cognate antigen during autoimmunity. Splenocytes from SFB-colonized control and IMQ-treated mice were restimulated ex vivo with the SFB peptide used for tetramer detection, and antigen recognition was assessed by Nur77 induction. Following SFB peptide restimulation, IMQ-treated mice exhibited increased numbers of Nur77⁺CD44⁺ CD4⁺ T cells, including Treg, RORγt⁺ Treg, Tfh, and Tfr populations (Fig. 2G; Fig. S3M-S). These findings indicate expansion of the SFB-reactive CD4⁺ T cell compartment during autoimmunity and are consistent with the broader accumulation of antigen-defined SFB tetramer⁺ T cells observed in vivo.

To determine whether these phenotypic and functional changes were accompanied by transcriptional remodeling, we performed bulk RNA sequencing of splenic SFB tetramer⁺ and tetramer-negative CD4⁺ T cells isolated from SFB-colonized IMQ-treated mice (Fig. S4A). Total splenic CD4⁺ T cells from IMQ-treated mice exhibited extensive transcriptional remodeling relative to healthy controls, with enrichment of proliferation, TCR signaling, type I interferon, and Th1/Tfh-associated pathways (Fig. S4B-E). Naïve-associated genes were suppressed, indicating a shift toward an activated effector state (Fig. S4D) (*19, 20*). Principal component analysis demonstrated clear segregation between SFB tetramer⁺ and tetramer-negative populations within IMQ-treated mice (Fig. S4B). Differential expression analysis identified enrichment of genes associated with T cell activation, proliferation, TCR signaling, and Th1/Tfh differentiation, including *Ifng, Il21, Maf, and Bhlhe40* in SFB tetramer⁺ T cells (Fig. S4F, G). Gene set enrichment analysis further confirmed the activation of Th1-, Tfh-, TCR signaling and activation-, and proliferation-associated pathways (Fig. 2H). These findings indicate that although autoimmunity broadly remodels the CD4⁺ T cell compartment, SFB tetramer⁺ T cells acquire a distinct transcriptional program enriched for activation, proliferation, and Th1/Tfh-associated pathways.

Together, these results demonstrate that TLR7-driven autoimmunity promotes the systemic accumulation of antigen-defined microbiota-reactive CD4⁺ T cells and drives their differentiation toward Th1- and Tfh-associated effector programs, as evidenced by coordinated phenotypic, functional, and transcriptional changes.

### Systemic autoimmunity promotes thymic generation of microbiota-reactive CD4⁺ T cells

Under homeostasis, thymic generation of microbiota-reactive CD4⁺ T cells is restricted to early life and is not induced by adult colonization (*16*). We therefore asked whether TLR7-driven autoimmunity could overcome this developmental restriction. Using I-Aᵇ/SFB3340 tetramers, we detected SFB tetramer⁺ T cells in the thymus of adult IMQ-treated SFB-colonized mice, while these cells remained undetectable following adult colonization under homeostatic conditions, indicating that systemic autoimmunity permits thymic accumulation of antigen-defined microbiota-reactive CD4⁺ T cells beyond the normal developmental window (Fig. 3A). As observed in the spleen, thymic accumulation was selective for SFB tetramer⁺ CD4⁺ T cells, whereas flagellin- and CLIP-specific CD4⁺ T cell frequencies were not significantly altered following IMQ treatment (Fig. S5A, B). Thymic SFB tetramer⁺ T cells also exhibited increased Ki67 expression compared with tetramer-negative CD4⁺ thymocytes, consistent with ongoing proliferation (Fig. 3B) (*18*). Systemic autoimmunity was accompanied by disruption of thymocyte development, characterized by reduced double-positive thymocytes and a relative enrichment of CD4⁺ and CD8⁺ single-positive populations (Fig. S5C-F). Alongside these global changes, SFB tetramer⁺ T cells accumulated within the thymus, suggesting preferential accumulation of microbiota-reactive populations during autoimmune-associated thymic remodeling.

**Fig. 3.**
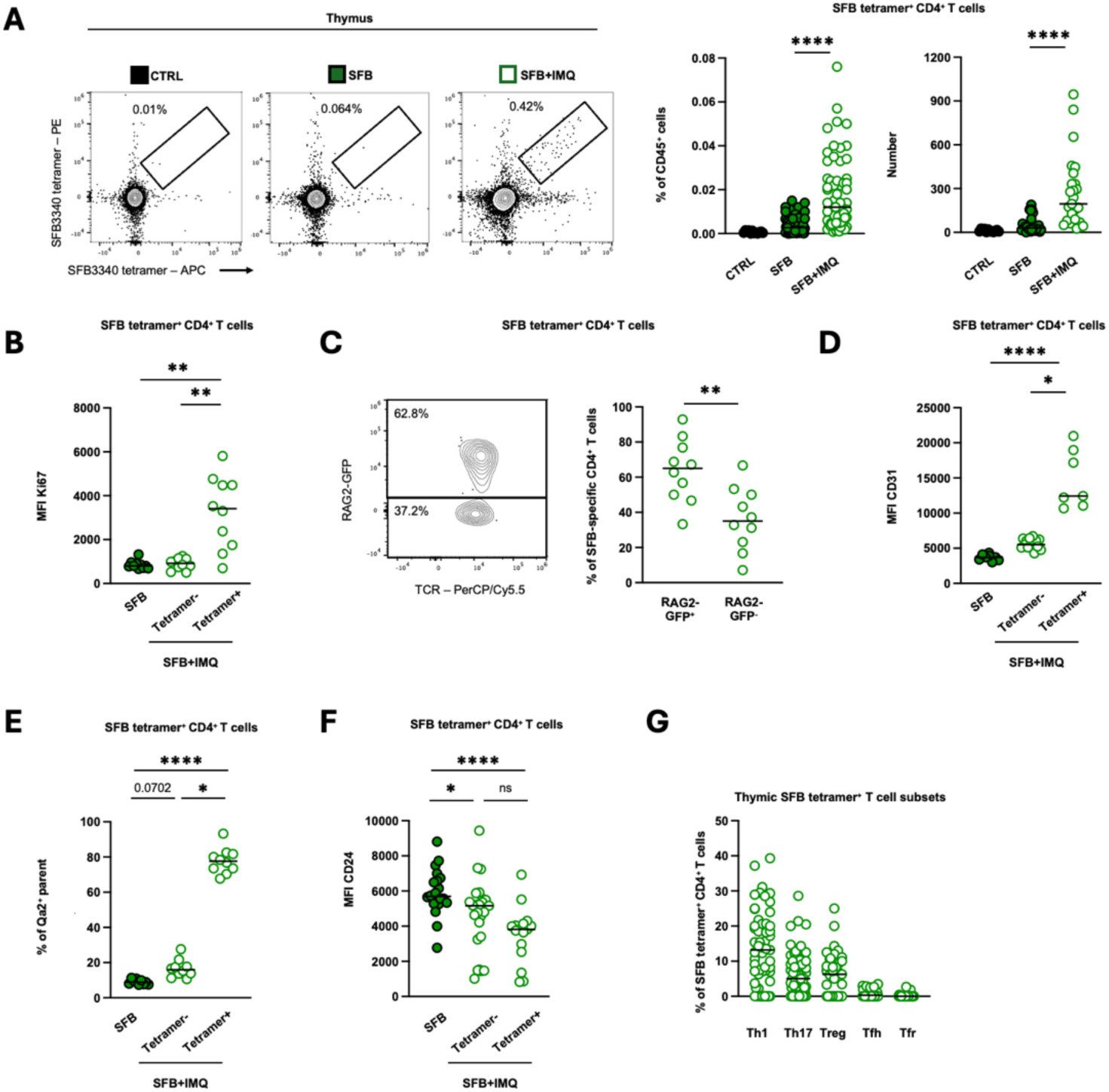
TLR7-driven autoimmunity overcomes developmental restriction of SFB tetramer⁺ CD4⁺ T cell generation. (A) SFB tetramer^+^ CD4⁺ T cells from thymus of control (CTRL), SFB-colonized (SFB), or SFB-colonized imiquimod-treated (SFB+IMQ) mice (n = 12-63). (B) Ki67 MFI in total CD4^+^ T cells from thymus of SFB and SFB tetramer^-^ or SFB tetramer^+^ CD4⁺ T cells from thymus of SFB+IMQ mice (n = 10). (C) GFP expression in SFB tetramer^+^ CD4⁺ T cells in thymus of reporter mice expressing GFP at the *rag2* locus (n = 10). (D-F) CD31 MFI (D), Qa2⁺ frequencies (E), and CD24 MFI (F) among thymic SFB tetramer^+^ CD4⁺ T cells (n = 8-25). (G) Th1, Th17, Treg, Tfh, and Tfr subsets in thymic SFB tetramer^+^ CD4⁺ T cells (n = 25-64). Data shown as means. Statistical significance was determined by Kruskal-Wallis test (A-B, D-F) or unpaired two-tailed Mann-Whitney U test (C). *\*p < 0.05, **p < 0.01, ***p < 0.001, ****p < 0.0001.* See also Figures S5 and S6.

Because peripheral T cells can recirculate to the thymus during inflammatory conditions (*12*), the presence of thymic SFB tetramer⁺ T cells does not by itself establish *de novo* thymic generation. To determine their developmental origin, we analyzed SFB tetramer⁺ T cells in RAG2-GFP reporter mice, in which recently generated thymocytes and recent thymic emigrants retain GFP expression following RAG2 expression during T cell development (*16*). We additionally assessed CD31 expression, another marker associated with recent thymic development (*21*). The majority of thymic SFB tetramer⁺ T cells expressed both RAG2-GFP and CD31 (Fig. 3C, D), consistent with newly generated thymocytes. The retention of both markers supports *de novo* thymic development rather than re-entry of mature peripheral T cells (*16*). Notably, a subset of splenic SFB tetramer⁺ T cells also retained RAG2-GFP and CD31 expression (Fig. S5G-I). IMQ-treated mice exhibited increased numbers of splenic RAG2-GFP⁺ SFB tetramer⁺ T cells, and splenic SFB tetramer⁺ T cells retained elevated CD31 expression, consistent with ongoing thymic export and peripheral accumulation of newly generated microbiota-reactive T cells during autoimmunity.

Despite promoting the generation of thymic SFB tetramer⁺ T cells, systemic autoimmunity was associated with a broader decline in thymic output. Frequencies of total RAG2-GFP⁺ CD4⁺ T cells were reduced in both the thymus and spleen of IMQ-treated mice (Fig. S5J, K), consistent with impaired thymopoiesis during chronic inflammation.

We next assessed the developmental status of thymic SFB tetramer⁺ T cells. Compared with tetramer-negative CD4⁺ thymocytes, SFB tetramer⁺ T cells exhibited increased Qa2 expression and reduced CD24 expression, consistent with a mature post-selection phenotype (Fig. 3E, F) (*22, 23*). Together, these findings indicate that newly generated microbiota-reactive CD4⁺ T cells acquire a mature post-selection phenotype within the adult thymus during systemic autoimmunity. In parallel, total thymic CD4⁺ T cells exhibited increased Qa2 expression and reduced CD24 expression compared with healthy controls (Fig. 3E, F), indicating altered thymocyte maturation during autoimmune-associated thymic remodeling. These findings indicate that microbiota-reactive CD4⁺ T cells are preferentially generated and undergo normal post-selection maturation despite a global reduction in thymic output during autoimmunity.

To determine whether newly generated thymic microbiota-reactive T cells already exhibited evidence of differentiation, we analyzed thymic SFB tetramer⁺ T cell subsets. Thymic SFB tetramer⁺ T cells exhibited a predominantly Th1 phenotype, with smaller Treg and Tfh populations also detected (Fig. 3G). Consistent with these findings, systemic autoimmunity increased Th1-, Treg-, and Tfh-associated populations within the broader thymic CD4⁺ T cell compartment (Fig. S5L). These results indicate that microbiota-reactive T cells begin acquiring features associated with peripheral effector differentiation during thymic development in autoimmune conditions. Consistent with their relatively early differentiation state, thymic SFB tetramer⁺ T cells produced minimal cytokines compared with their splenic counterparts, indicating that effector programming is initiated in the thymus but becomes fully established following peripheral expansion (Fig. S5M-Q).

To determine whether these phenotypic changes were accompanied by transcriptional remodeling, we performed bulk RNA sequencing of thymic SFB tetramer⁺ and tetramer-negative CD4⁺ T cells isolated from SFB-colonized IMQ-treated mice. Compared with tetramer-negative thymocytes, SFB tetramer⁺ T cells upregulated the expression of genes associated with proliferation and TCR signaling, while suppressing naïve-associated programs (Fig. S6A-D). Although the magnitude of transcriptional remodeling was more modest than that observed in splenic SFB tetramer⁺ T cells, many of the genes within these pathways overlapped between tissues, including programs associated with proliferation, activation, and Th1/Tfh differentiation. These findings indicate that microbiota-reactive CD4⁺ T cells begin acquiring effector-associated transcriptional features during thymic development (*20*), which are subsequently reinforced following peripheral expansion during systemic autoimmunity.

Together, these findings demonstrate that TLR7-driven autoimmunity overcomes the developmental restriction that normally limits thymic generation of microbiota-reactive CD4⁺ T cells in adulthood. Newly generated SFB tetramer⁺ T cells emerge in the thymus, acquire early Th1/Tfh-associated transcriptional and phenotypic features, and contribute to the pool of microbiota-reactive T cells that accumulates in peripheral tissues during systemic autoimmunity.

### Microbiota-reactive CD4⁺ T cells emerge early and accumulate systemically during spontaneous lupus

To determine whether the accumulation of microbiota-reactive T cells observed in inducible lupus also occurs during spontaneous disease, we analyzed SFB tetramer⁺ T cells in TLR7.1 transgenic (TLR7.1 Tg) mice, which spontaneously develop lupus-like disease and display overt clinical manifestations, including kidney involvement, by 12 weeks of age, driven by TLR7 overexpression (*17*). In contrast to the milder pathology seen in IMQ-treated mice at 6 weeks of treatment, this model allows assessment of microbiota-reactive T cells across disease stages. To investigate microbiota-reactive T cells throughout disease progression, we analyzed SFB tetramer⁺ T cells at two stages of disease: 5 weeks of age (young), following colonization at weaning and before overt disease, and 14 weeks of age (adult), following colonization at 12 weeks and after severe systemic pathology had developed (Fig. 4A) (*17*).

**Fig. 4.**
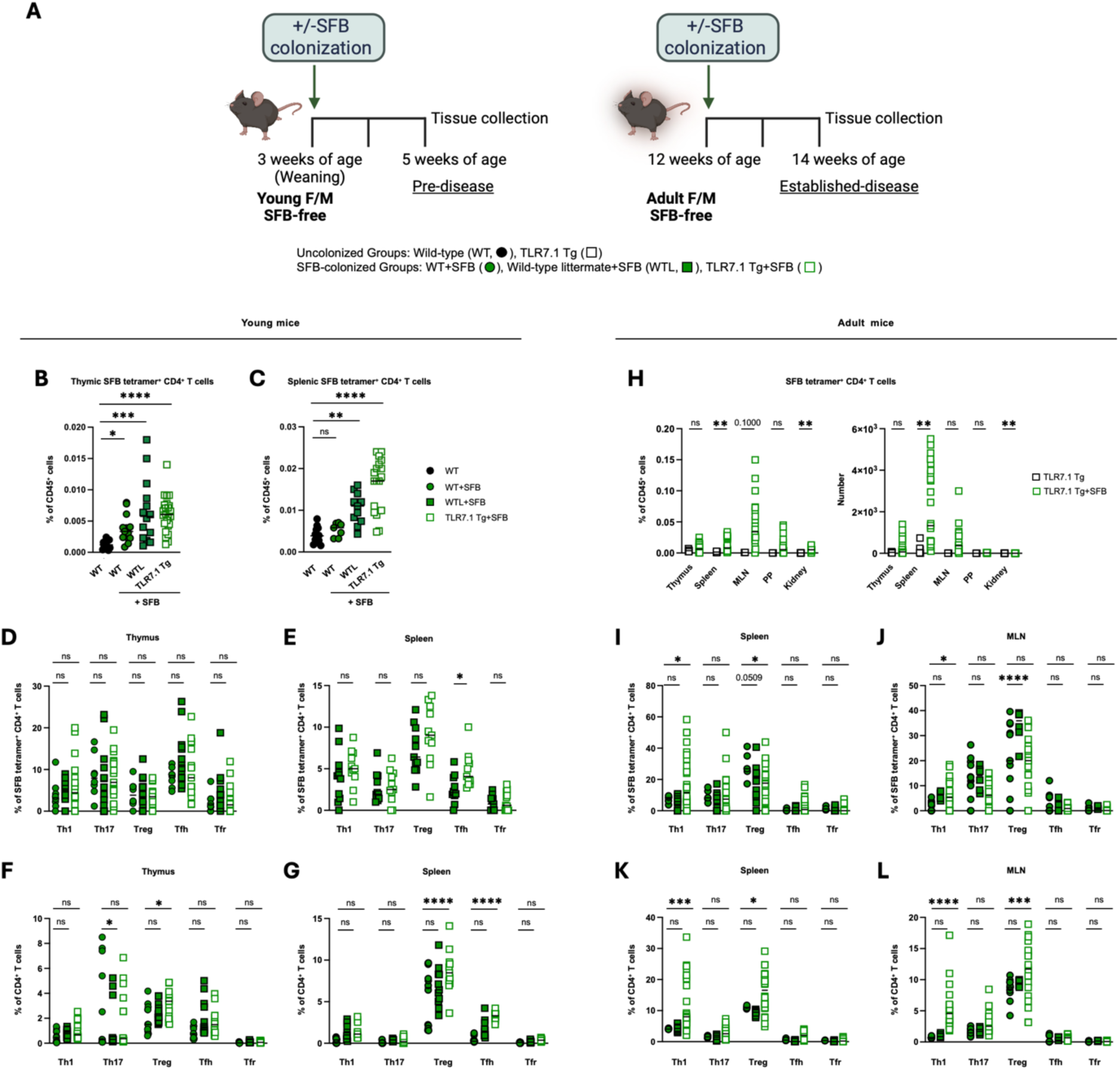
SFB tetramer⁺ CD4⁺ T cells emerge early and accumulate during disease progression in TLR7-driven autoimmunity. (A) Experimental design for evaluating SFB tetramer^+^ CD4⁺ T cell responses in thymus and spleen of wild-type (WT) mice, WT littermates (WTL), and TLR7.1 Tg mice, two weeks after SFB colonization. Young mice were colonized at three weeks and tissues collected at five weeks, and adult mice were colonized at 12 weeks and tissues collected at 14 weeks. (B and C) Thymic (B) and splenic (C) SFB tetramer^+^ CD4⁺ T cells (n = 7-24). (D and E) Effector subsets of SFB tetramer^+^ CD4^+^ T cells in thymus (D) and spleen (E) of young mice (n = 8-17). (F and G) Effector subsets of total CD4^+^ T cells in thymus (F) and spleen (G) of young mice (n = 8-19). (H) SFB tetramer^+^ CD4^+^ T cells across tissues (n = 2-27). (I and J) SFB tetramer^+^ CD4^+^ T cell effector subsets in spleen (I) and mesenteric lymph node (MLN) of adult mice (n = 5-28). (K and L) Total CD4⁺ T cell subsets in spleen (K) and MLN (L) of adult mice (n = 5-28). Panel A was created with Biorender.com. Data shown as means. Statistical significance was determined by Kruskal-Wallis test (B, C), two-way ANOVA (D, F-G, I-L), and multiple unpaired two-tailed Mann-Whitney U tests (E, H). *\*p < 0.05, **p < 0.01, ***p < 0.001*.

In young TLR7.1 Tg mice analyzed 2 weeks after weaning, SFB tetramer⁺ T cells were detected in both the thymus and spleen (Fig. 4B, C). While thymic SFB tetramer⁺ T cells are normally generated during early life (*16*), their early appearance in peripheral tissues was not observed in young healthy wild-type controls. These findings suggest that lupus predisposition promotes the early peripheral emergence of microbiota-reactive T cells prior to the development of overt disease. Notably, wild-type littermate (WTL) controls with normal TLR7 expression cohoused with TLR7.1 Tg mice also exhibited increased splenic SFB tetramer⁺ T cells relative to healthy controls (Fig. 4B, C). Phenotypic analysis revealed limited differentiation of thymic SFB tetramer⁺ T cells at this early stage, whereas splenic SFB tetramer⁺ T cells exhibited increased Tfh polarization in TLR7.1 Tg mice (Fig. 4D, E). Similar changes were observed within the total CD4⁺ T cell compartment, where both TLR7.1 Tg and WTL mice exhibited increased Tfh frequencies in the thymus and spleen, while TLR7.1 Tg mice additionally showed increased Treg frequencies in both tissues (Fig. 4F, G). These findings suggest that microbiota-reactive T cells undergo peripheral differentiation early during lupus predisposition within a broader program of immune remodeling that precedes overt disease.

In adult TLR7.1 Tg mice with established disease (*9*), SFB tetramer⁺ T cells were detected across multiple tissues, including the thymus, spleen, MLN, Peyer’s patches, and kidney, in SFB-colonized mice, indicating systemic dissemination of microbiota-reactive T cells in established disease (Fig. 4H). Although SFB tetramer⁺ T cells were most abundant in secondary lymphoid and gut-associated tissues, they were also detected at low frequencies in the kidney, indicating access to a major target organ of lupus-associated inflammation. We next examined whether microbiota-reactive T cells exhibited altered differentiation during established disease. Comparison with wild-type and wild-type littermate controls revealed reciprocal enrichment of Th1 cells and reduction of Tregs among SFB tetramer⁺ T cells in both the spleen and MLN of TLR7.1 Tg mice (Fig. 4I, J), consistent with inflammatory reprogramming of microbiota-reactive T cells during lupus progression. Similar remodeling was observed within the total CD4⁺ T cell compartment, which exhibited increased Th1 and Treg frequencies in both tissues (Fig. 4K, L). Together, these findings demonstrate that microbiota-reactive T cells emerge early during lupus predisposition and undergo widespread dissemination and inflammatory reprogramming as disease progresses. While early responses are characterized by increased Tfh differentiation, established disease is associated with systemic accumulation of microbiota-reactive T cells and inflammatory reprogramming toward Th1-biased effector states.

### Intestinal dendritic cell trafficking promotes microbiota-reactive CD4⁺ T cell accumulation during autoimmunity

To investigate how systemic autoimmunity promotes the accumulation of microbiota-reactive CD4⁺ T cells, we examined dendritic cell (DC) trafficking in SFB-colonized mice with or without IMQ treatment as described above. Under homeostatic conditions, we previously identified a population of intestinal CX3CR1⁺ antigen-presenting cells (APCs) that migrate to the thymus and promote the development of microbiota-reactive CD4⁺ T cells (*16*). We therefore hypothesized that systemic autoimmunity reactivates this trafficking pathway in adult mice.

Analysis of dendritic cell populations revealed that IMQ-treated mice exhibited an increased frequency of total DCs in the thymus, without a corresponding increase in absolute numbers (Fig. S7A, B), consistent with inflammation-associated thymic involution (*24, 25*). In contrast, splenic DC numbers increased substantially during autoimmunity (Fig. S7C). Within this broader remodeling, we observed a significant increase in the frequency and number of CD11b⁺ DCs in both the thymus and spleen of IMQ-treated mice (Fig. 5A, B). These cells expressed SIRPα, CX3CR1, CCR7, and IRF4, lacked expression of macrophage-associated markers, and were negative for XCR1, CD103, and IRF8 (Fig. S7D-F). Importantly, expansion of CD11b⁺ DCs was not accompanied by widespread alterations in other antigen-presenting cell populations (Fig. S7G). Although other splenic APC subsets exhibited modest increases in frequency, the most prominent and consistent APC alteration observed during autoimmunity was the expansion of CD11b⁺ DCs, which displayed phenotypic features highly consistent with migratory cDC2s (*26, 27*). This population closely resembles the gut-derived APC subset previously shown to traffic to the thymus and promote the development of microbiota-reactive CD4⁺ T cells under homeostatic conditions *(16)*. These findings identify an expanded CD11b⁺ dendritic cell population in the thymus and spleen during systemic autoimmunity.

**Fig. 5.**
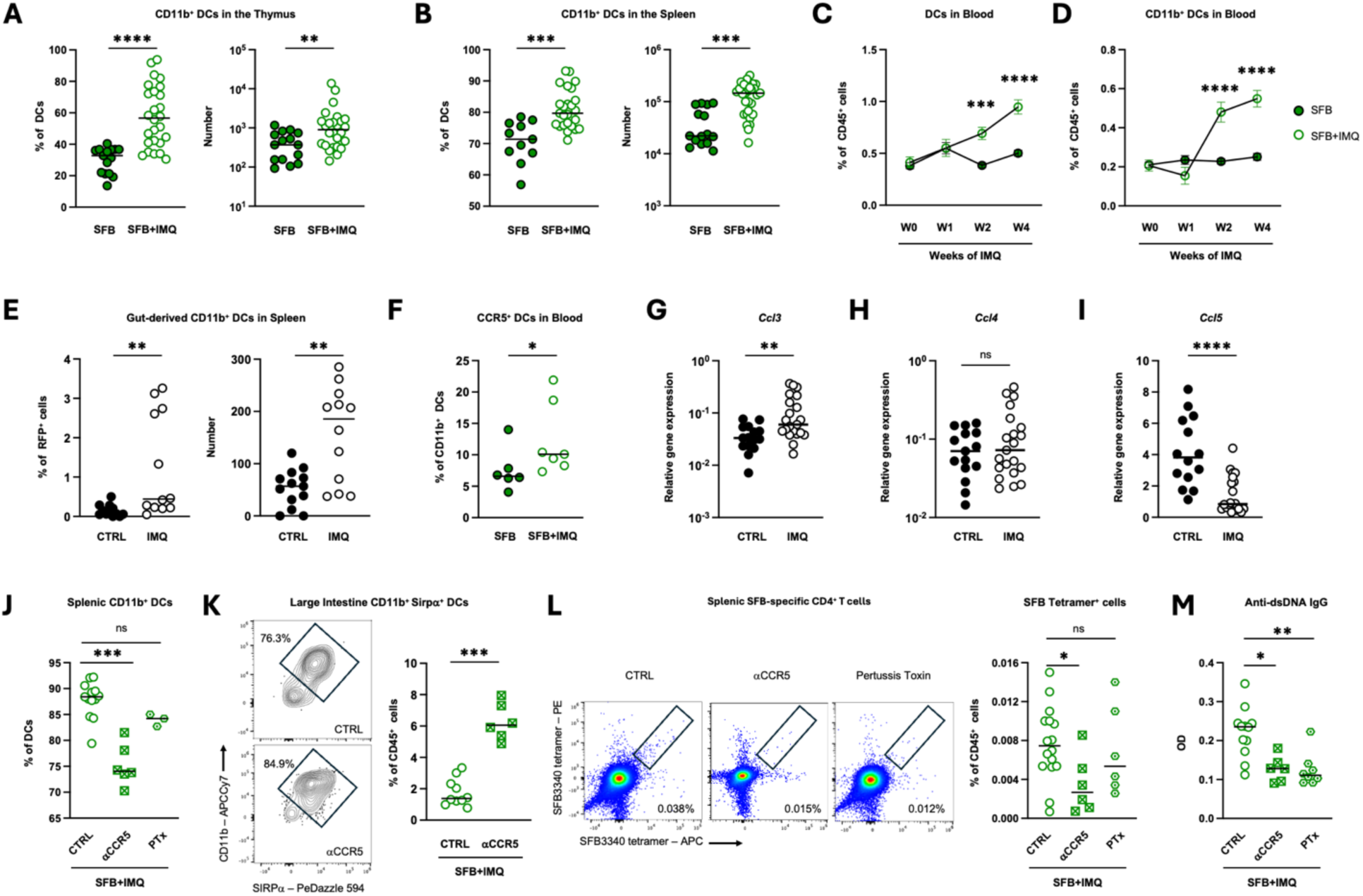
Reactivation of intestinal dendritic cell trafficking supports SFB tetramer⁺ CD4⁺ T cell expansion during TLR7-driven autoimmunity. (A and B) Thymic (A) and splenic (B) CD11b⁺ DC cell populations in SFB and SFB+IMQ mice (n = 11-29). (C and D) Circulating total and CD11b⁺ DCs from SFB-colonized mice with or without 0 to 4 weeks of imiquimod (IMQ) treatment (n = 15-18). (E) Splenic RFP⁺ CD11b⁺ DCs post-cecal photoconversion in KikGR reporter mice (n = 5). (F) Frequency of CCR5-expressing CD11b^+^ DCs in blood (n = 6-7). (G-I) Gene expression of *Ccl3* (G), *Ccl4* (H), and *Ccl5* (I) in spleen (n = 14-21). (J) Frequency of CD11b^+^ cells of total DCs in spleen after anti-CCR5 blockade (αCCR5) or pertussis toxin (PTx) treatment (n = 3-13). (K) Frequency of CD11b^+^ SIRPα^+^ DCs in large intestine after αCCR5 blockade (n = 6-10) (L-M) SFB tetramer^+^ CD4⁺ T cells in spleen (L) and serum anti-dsDNA IgG (M) after αCCR5 or PTx treatment (n = 6-16). Data shown as means. Statistical significance was determined by Kruskal-Wallis test (J, L-M), two-way ANOVA (C-D), and unpaired two-tailed Mann-Whitney U test (A-B, E-I, K). *\*p < 0.05, **p < 0.01, ***p < 0.001, ****p < 0.0001.* See also Figure S7.

We next asked whether intestinal DC trafficking was increased during systemic autoimmunity. Longitudinal blood analysis revealed a progressive rise in circulating DCs throughout IMQ treatment, particularly within the CD11b⁺ compartment (Fig. 5C, D). Although transient changes were observed in several circulating immune populations early after IMQ initiation, including neutrophils, monocytes, B cells, and T cells, these alterations largely normalized over time (Fig. S7H-L). In contrast, circulating DCs remained persistently elevated throughout disease progression, identifying sustained DC expansion as a distinguishing feature of systemic autoimmunity. We therefore sought to determine whether the expanded CD11b⁺ DC population originated from the intestine and reflected enhanced intestinal DC migration. To address this, we used photoconvertible KikGR mice and selectively exposed the cecum to 405-nm light, converting local GFP⁺ cells to RFP⁺ (*28*). Twenty-four hours later, photoconverted intestinal CD11b⁺ DCs were readily detected in the spleen, with significantly greater frequencies and numbers in IMQ-treated mice than in controls (Fig. 5E). These findings demonstrate that systemic autoimmunity enhances the dissemination of intestinal migratory CD11b⁺ DCs to peripheral lymphoid tissues (*16, 29, 30*).

Because intestinal CX3CR1⁺ APCs require CCR5-dependent trafficking to access the thymus and support microbiota-reactive CD4⁺ T cell development under homeostatic conditions (*16*), we next tested whether this pathway contributes to microbiota-reactive CD4⁺ T cell accumulation during autoimmunity. Consistent with altered CCR5-associated signaling during disease, circulating CD11b⁺ DCs from IMQ-treated mice expressed higher levels of CCR5 than controls (Fig. 5F). In parallel, expression of CCR5 ligands was altered in the spleen, characterized by increased *ccl3*, unchanged *ccl4*, and decreased *ccl5* expression (Fig. 5G-I). These findings suggested involvement of the CCR5 axis during autoimmunity and prompted us to test whether CCR5 blockade could limit the accumulation of microbiota-reactive CD4⁺ T cells during disease. Mice therefore received a CCR5 antagonist during the final two weeks of IMQ treatment (*31*). CCR5 blockade reduced the accumulation of CD11b⁺ DCs in the spleen while increasing their retention within the intestine, consistent with impaired intestinal DC trafficking (Fig. 5J, K). This intervention significantly reduced the frequency of splenic SFB tetramer⁺ T cells, accompanied by a reduction in total splenic CD4⁺ T cell frequencies, while preserving the Th1 and Tfh composition of the remaining SFB tetramer⁺ T cell population (Fig. 5L; Fig. S7M, N). CCR5 blockade also significantly reduced anti-dsDNA autoantibody levels, while causing only modest effects on spleen size and anti-ssRNA autoantibody responses (Fig. 5M, S7O, P). Together, these findings identify CCR5-dependent intestinal DC trafficking as an important regulator of microbiota-reactive CD4⁺ T cell accumulation during lupus. To further examine the contribution of chemokine-dependent immune cell trafficking, mice received pertussis toxin (PTx), an inhibitor of Gi-coupled chemokine receptor signaling, during the final two weeks of IMQ treatment. PTx treatment did not significantly alter splenic CD11b⁺ DC frequencies, SFB tetramer⁺ T cell accumulation, total CD4⁺ T cell numbers, or tetramer⁺ T cell subset composition (Fig. 5J, L; Fig. S7M, N). Despite these limited effects on cellular populations, PTx significantly reduced both anti-dsDNA and anti-ssRNA autoantibody levels without affecting splenomegaly (Fig. 5M, S7O, P). These findings suggest that broader disruption of migration-dependent pathways can attenuate humoral autoimmune responses in lupus, although the cellular mechanisms underlying these effects differ from those observed following CCR5 blockade.

Together, these findings demonstrate that systemic autoimmunity enhances intestinal dendritic cell trafficking and identify CCR5-dependent migration as an important regulator of microbiota-reactive CD4⁺ T cell accumulation during lupus. More broadly, perturbation of immune cell trafficking pathways attenuated selected manifestations of autoimmune pathology. These results identify intestinal immune migration as a mechanistic link between the gut microbiota and systemic autoimmunity.

### TLR7-driven autoimmunity amplifies pathobiont-reactive CD4⁺ T cell responses

Having established that TLR7-driven autoimmunity promotes the expansion and differentiation of microbiota-reactive CD4⁺ T cells, we next asked how autoimmunity influences pathobiont-reactive responses. *E. gallinarum* (EG) has been linked to lupus pathogenesis in both mice and humans and exacerbates disease in TLR7-driven autoimmunity (*4, 5*). We therefore examined how TLR7-dependent autoimmunity shapes *E. gallinarum*-reactive CD4⁺ T cell responses (Fig. 6A).

**Fig. 6.**
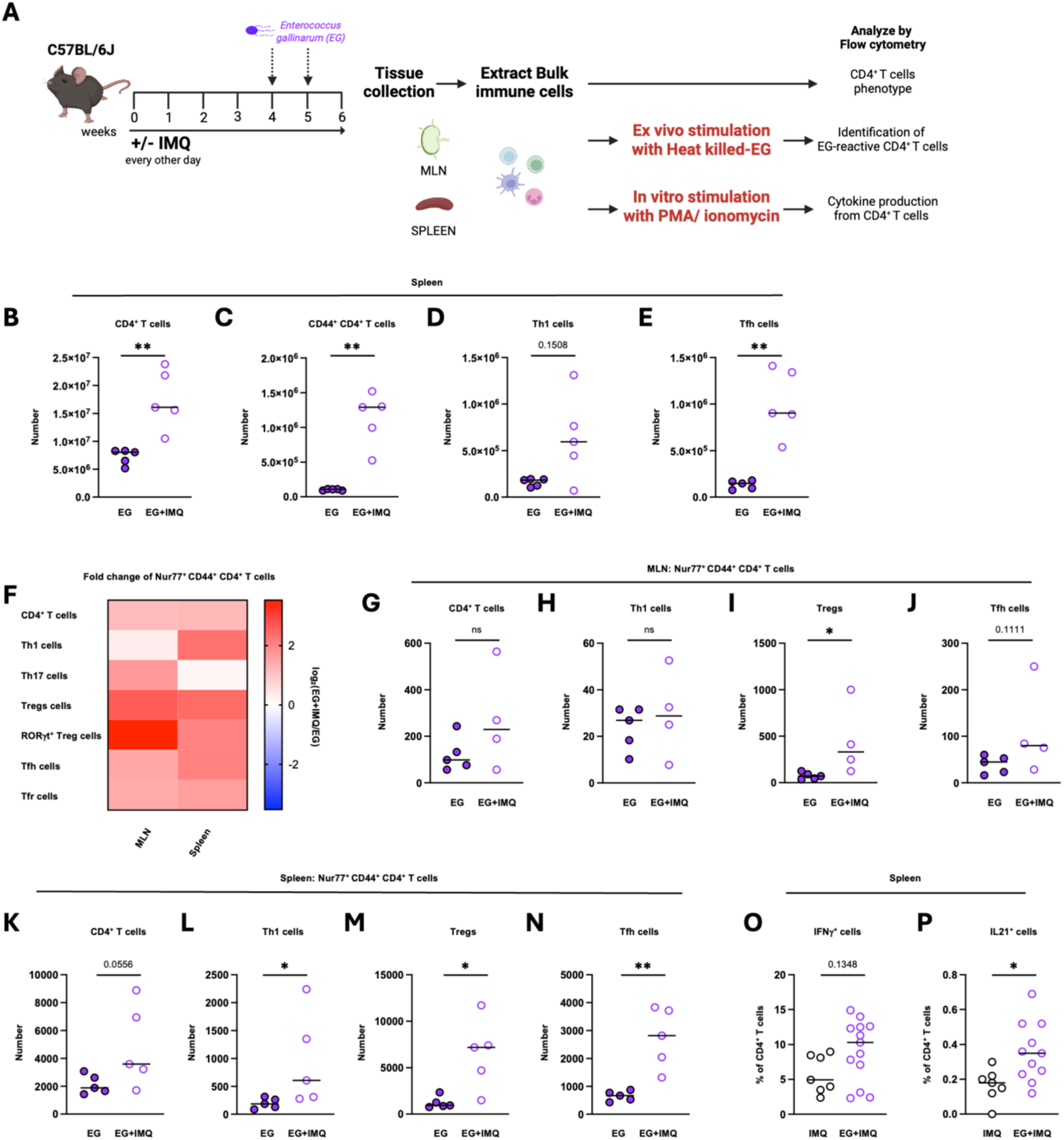
TLR7-driven autoimmunity promotes Th1- and Tfh-associated pathobiont-reactive CD4⁺ T cell responses. (A) Experimental schematic of in vitro T cell stimulation with PMA/Ionomycin and ex vivo T cell restimulation with heat-killed EG. (B-E) Counts of total (B), CD44⁺ (C), Th1 (D), or Tfh (E) CD4^+^ T cells in spleen of *E. gallinarum*-colonized control (EG) and EG-colonized IMQ-treated (EG+IMQ) mice (n = 5). (F) Fold change of Nur77⁺ cells among CD4⁺ T cells and their subsets in the MLN and spleen of *E. gallinarum*-colonized control (EG) and EG-colonized IMQ-treated (EG+IMQ) mice (n = 5). (G-N) Counts of total CD4^+^ (G, K), Th1 (H, L), Treg (I, M), and Tfh (J, N) Nur77^+^ cells in the MLN (G-J) and spleen (K-N) of IMQ and EG+IMQ mice (n = 4-5). (O-P) Frequency of IFNγ (O) and IL-21 (P) producing cells among CD4⁺ T cells from IMQ and EG+IMQ mice (n = 7-13). Panel A was created with Biorender.com. Data shown as means. Statistical significance was determined by unpaired two-tailed Mann-Whitney U test (B-E, G-P). *\*p < 0.05, **p < 0.01, ***p < 0.001, ****p < 0.0001*.

Compared with mice colonized with *E. gallinarum* alone, EG-colonized IMQ-treated mice exhibited increased numbers of splenic CD4⁺ T cells and a marked expansion of the CD44⁺ antigen-experienced compartment (Fig. 6B, C). This expansion was accompanied by substantial remodeling of CD4⁺ T cell subsets, including increased Th1 and Tfh populations, consistent with enhanced differentiation during autoimmunity (Fig. 6D, E). These findings indicate that TLR7-driven autoimmunity promotes broad remodeling of CD4⁺ T cell responses in *E. gallinarum*-colonized mice.

We next assessed pathobiont-reactive responses using ex vivo restimulation with *E. gallinarum* lysates. Compared with untreated *E. gallinarum*-colonized mice, IMQ-treated mice exhibited increased *E. gallinarum*-reactive CD44⁺ CD4⁺ T cell responses in both the MLN and spleen (Fig. 6F). In the MLN, enhanced responses were observed across multiple CD4⁺ T cell subsets, indicating that systemic autoimmunity amplifies pathobiont-reactive T cell responses within gut-draining lymphoid tissues. In the spleen, increased numbers of antigen-reactive CD4⁺ T cell subsets, particularly Th1- and Tfh-associated populations, were detected following *E. gallinarum* restimulation (Fig. 6G-N). These findings indicate that systemic autoimmunity amplifies pathobiont-reactive CD4⁺ T cell responses and promotes the accumulation of pathobiont-reactive Th1- and Tfh-associated populations.

To determine whether *E. gallinarum* exposure further altered effector function during autoimmunity, we examined cytokine production by splenic CD4⁺ T cells. Compared with IMQ-treated mice, EG-colonized IMQ-treated mice exhibited increased frequencies of IFNγ⁺ and IL-21⁺ CD4⁺ T cells, consistent with enhanced Th1 and Tfh effector programs (Fig. 6O, P) (*32, 33*). These findings indicate that *E. gallinarum* amplifies Th1- and Tfh-associated responses during systemic autoimmunity.

Together, these findings demonstrate that TLR7-driven autoimmunity promotes the expansion and differentiation of pathobiont-reactive CD4⁺ T cells, while *E. gallinarum* further enhances Th1- and Tfh-associated programs. These observations support the possibility that pathobiont-reactive CD4⁺ T cells contribute directly to disease progression.

### Pathobiont-conditioned CD4⁺ T cells exacerbate autoimmunity through IL-21- and IFNγ-associated pathways

Having established that TLR7-driven autoimmunity promotes the expansion and differentiation of *E. gallinarum*-reactive CD4⁺ T cells, and that *E. gallinarum* further enhances Th1- and Tfh-associated effector programs, we next asked whether these cells directly contribute to disease progression. Splenic CD4⁺ T cells were isolated from SFB-colonized, EG-colonized, or uncolonized IMQ-treated donors and transferred into colonized RAG2-deficient recipients together with B cells from uncolonized donors. Recipient mice were subsequently treated with IMQ for six weeks. To determine the contribution of IL-21 and IFNγ signaling to *E. gallinarum*-driven pathology, a subset of recipient mice received IL-21R-blocking antibodies alone or in combination with IFNγ blockade throughout disease progression (Fig. 7A).

**Fig. 7:**
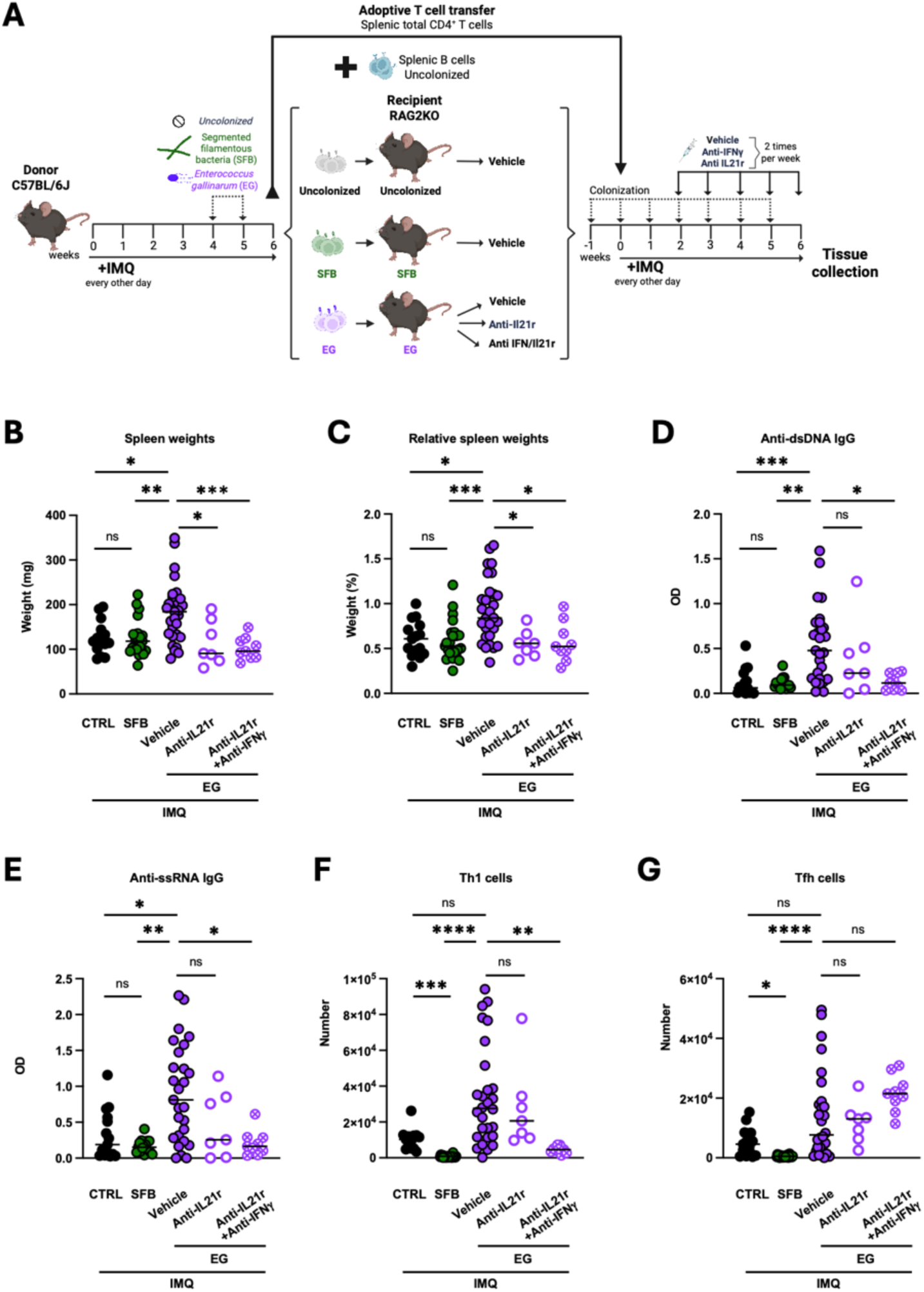
Pathobiont-reactive CD4^+^ T cells exacerbate autoimmunity through IL-21 and IFNγ-associated pathways. (A) Experimental schematic of adoptive transfer of splenic CD4⁺ T cells. IMQ-treated wild-type mice were left uncolonized (CTRL) or were colonized with either segmented filamentous bacteria (SFB) or *Enterococcus gallinarum* (EG). Recipient RAG2KO mice were colonized to match donor status, co-transferred with B cells from uncolonized mice, and treated with IMQ for 6 weeks after transfer. To assess cytokine contributions to disease exacerbation, some recipients received anti-IL-21 with or without concomitant anti-IFNγ blockade from weeks two to six of IMQ treatment. (B and C) Spleen weights (B) and spleen/body weight ratio (C) (n = 7-29). (D and E) Serum anti-dsDNA (D) and anti-ssRNA (E) IgG (n = 7-27). (F-G) Counts of splenic Th1 (F), and Tfh (G) CD4^+^ T cells (n = 6-21). Panel A was created with Biorender.com. Data shown as means. Statistical significance was determined by Kruskal-Wallis (B-G). *\*p < 0.05, **p < 0.01, ***p < 0.001, ****p < 0.0001.* See also Figures S8 and S9.

Transfer of CD4⁺ T cells from SFB-colonized donors did not alter splenomegaly, autoantibody production, or splenic effector T cell accumulation compared with control donor cells. In contrast, recipients of *E. gallinarum*-conditioned CD4⁺ T cells developed substantially greater splenomegaly, elevated anti-dsDNA and anti-ssRNA autoantibody levels, and enhanced Th1 and Tfh responses (Fig. 7B-G). These findings demonstrate that *E. gallinarum*-conditioned CD4⁺ T cells exhibit greater pathogenic potential than CD4⁺ T cells from SFB-colonized donors. Compared with recipients of *E. gallinarum*-conditioned CD4⁺ T cells that did not receive cytokine blockade, IL-21R blockade alone modestly reduced splenomegaly but did not significantly alter autoantibody production or splenic Th1 and Tfh populations (Fig. 7B-G). In contrast, combined IL-21R and IFNγ blockade markedly reduced disease severity, including splenomegaly, anti-dsDNA and anti-ssRNA IgG responses, and Th1 cell accumulation. These findings indicate that IL-21R blockade alone is insufficient to fully reverse *E. gallinarum*-mediated pathology, whereas combined IL-21R and IFNγ blockade substantially attenuates disease severity, supporting a role for both cytokine pathways in exacerbation of pathobiont-driven disease. Although both commensal- and pathobiont-reactive CD4⁺ T cells acquired effector phenotypes during autoimmunity (Figs. 2 and 6), only *E. gallinarum*-conditioned CD4⁺ T cells exhibited substantial pathogenic activity following adoptive transfer, markedly amplifying autoantibody production and effector T cell accumulation. These findings demonstrate that microbial context is a critical determinant of microbiota-reactive CD4⁺ T cell pathogenicity and identify IL-21- and IFNγ-dependent pathways as important mediators of pathobiont-driven disease exacerbation.

To determine whether thymic CD4⁺ T cells generated following *E. gallinarum* colonization could also contribute to disease, we performed adoptive transfer experiments using thymic CD4⁺ T cells isolated from EG-colonized and uncolonized IMQ-treated donors. RAG2KO recipients were colonized with *E. gallinarum*, co-transferred with B cells from uncolonized IMQ-treated donors, and subsequently treated with IMQ for six weeks (Fig. S8A). For both groups, recipients of thymic CD4⁺ T cells developed only modest disease manifestations. *E. gallinarum*-colonized recipients exhibited mild splenomegaly (Fig. S8B, C), while anti-dsDNA and anti-ssRNA IgG levels showed upward trends that did not reach statistical significance (Fig. S8D, E). Despite these changes, transfer of thymic CD4⁺ T cells failed to significantly alter splenic effector T cell populations (Fig. S8F, G). These findings indicate that thymic *E. gallinarum*-reactive CD4⁺ T cells possess limited pathogenic activity and suggest that peripheral conditioning is required for the full disease-promoting potential of pathobiont-reactive CD4⁺ T cells.

We next asked whether *E. gallinarum*-conditioned CD4⁺ T cells could promote disease in the presence of an intact immune system and ongoing pathobiont exposure. To address this, splenic CD4⁺ T cells from IMQ-treated CD45.1⁺ donor mice colonized with *E. gallinarum* or left uncolonized were transferred into IMQ-treated immunocompetent CD45.2⁺ recipients previously colonized with *E. gallinarum* (Fig. S9A). *E. gallinarum*-conditioned CD4⁺ T cells retained pathogenic activity despite transfer into immunocompetent hosts. Recipients of *E. gallinarum*-conditioned donor cells developed splenomegaly and elevated anti-dsDNA and anti-ssRNA autoantibody levels compared with recipients of control donor cells (Fig. S9B-E). In addition, *E. gallinarum*-conditioned CD4⁺ T cells promoted a strong Tfh bias within endogenous CD45.2⁺ CD4⁺ T cell populations, indicating their capacity to reshape ongoing immune responses within an intact host (Fig. S9F, G). These findings demonstrate that *E. gallinarum*-conditioned CD4⁺ T cells retain pathogenic activity even in immunocompetent hosts with ongoing *E. gallinarum* exposure and established autoimmune inflammation. Their ability to amplify disease and reshape endogenous CD4⁺ T cell responses indicates that pathobiont-conditioned T cells can actively propagate, rather than merely initiate, autoimmune pathology.

Collectively, these results demonstrate that *E. gallinarum*-conditioned CD4⁺ T cells amplify autoimmunity through IL-21- and IFNγ-dependent mechanisms. In direct comparison with CD4⁺ T cells from SFB-colonized donors, *E. gallinarum*-conditioned CD4⁺ T cells exhibited substantially greater pathogenic potential despite inducing similar effector phenotypes. *E. gallinarum*-conditioned CD4⁺ T cells promoted a pathogenic program characterized by enhanced Tfh differentiation, autoantibody production, and systemic immune activation, which could be reversed by cytokine blockade. These findings establish that microbial context is a critical determinant of microbiota-reactive CD4⁺ T cell pathogenicity and identify IL-21- and IFNγ-dependent pathways as key mediators of pathobiont-driven disease exacerbation.

## DISCUSSION

Our study demonstrates that systemic autoimmunity broadly reshapes microbiota-reactive CD4⁺ T cell immunity and that microbial identity determines the pathogenic consequences of these responses. We found that systemic autoimmunity enhanced microbiota-reactive CD4⁺ T cell responses, promoted the accumulation and differentiation of antigen-defined microbiota-reactive T cells, and increased their presence across lymphoid and peripheral tissues. Despite acquiring Th1 and Tfh characteristics during disease, commensal-reactive CD4⁺ T cells exhibited limited pathogenic potential following adoptive transfer. In contrast, pathobiont-conditioned CD4⁺ T cells exhibited substantially greater pathogenic potential and exacerbated autoimmune pathology through distinct cytokine-dependent mechanisms. Together, these findings support a model in which systemic autoimmunity reshapes microbiota-reactive CD4⁺ T cell immunity, while microbial context determines whether these responses acquire pathogenic potential.

Microbiota-reactive CD4⁺ T cells are abundant components of the normal immune repertoire and contribute to immune homeostasis and host defense at barrier surfaces. Previous studies in humans demonstrated that these cells undergo functional alterations during chronic inflammatory disease, suggesting that inflammatory environments can reshape microbiota-reactive immunity (*12*). Our findings extend these observations by showing that systemic autoimmunity broadly remodels microbiota-reactive CD4⁺ T cell responses across intestinal and systemic lymphoid tissues. Using fecal lysate stimulation and antigen-defined SFB tetramer⁺ T cells, we found that autoimmunity promotes the systemic accumulation and differentiation of microbiota-reactive CD4⁺ T cells. Transcriptomic, phenotypic, and functional analyses collectively revealed enrichment of Th1 and Tfh programs, indicating that autoimmunity alters not only the magnitude but also the quality of microbiota-reactive T cell responses. Notably, despite previous reports linking SFB and lupus-associated pathobionts to Th17 responses (*4, 5, 26, 34*), microbiota-reactive CD4⁺ T cells in our models predominantly acquired Th1 and Tfh characteristics, mirroring the broader polarization of the total CD4⁺ T cell compartment during systemic autoimmunity. Together, these findings identify autoimmune inflammation as a potent regulator of microbiota-reactive CD4⁺ T cell differentiation and suggest that disease-associated immune environments can reprogram host–microbiota interactions by promoting effector programs associated with autoimmunity.

Our findings further suggest that altered intestinal immune trafficking contributes to the accumulation of microbiota-reactive CD4⁺ T cells during systemic autoimmunity. We observed increased migration of gut-derived dendritic cells to both the spleen and thymus during disease, accompanied by expansion of microbiota-reactive T cells, which was reduced following perturbation of trafficking pathways. These observations extend our previous work demonstrating that intestinal CX3CR1⁺ antigen-presenting cells support microbiota-specific T cell development during early life and suggest that autoimmunity reactivates components of this developmental circuit in adulthood (*16*). Notably, these findings indicate that systemic autoimmunity overcomes the developmental restriction normally imposed on the generation of microbiota-reactive T cells, permitting their emergence beyond the early-life window observed under homeostatic conditions. Consistent with this model, microbiota-reactive CD4⁺ T cells were detected in the thymus of adult lupus models, while analysis of TLR7.1 transgenic mice revealed early peripheral accumulation and progressive systemic expansion of these populations during disease progression. Although the precise mechanisms governing intestinal dendritic cell egress and tissue targeting remain unclear, our data support a model in which inflammatory signals promote the dissemination of gut-derived immune cells beyond the intestine, thereby driving the systemic accumulation and peripheral differentiation of microbiota-reactive CD4⁺ T cells during autoimmunity (Fig. S10).

Despite undergoing substantial expansion and differentiation during autoimmunity, not all microbiota-reactive CD4⁺ T cells exhibited equivalent pathogenic potential. SFB tetramer⁺ T cells accumulated systemically and adopted Th1 and Tfh characteristics, but exhibited limited pathogenic activity following adoptive transfer. In contrast, *Enterococcus gallinarum*-conditioned CD4⁺ T cells amplified autoimmune pathology through IL-21- and IFNγ-dependent effector programs to exacerbate disease. Notably, these pathogenic effects persisted after transfer into immunocompetent recipients, indicating that pathobiont-conditioned CD4⁺ T cells can reshape ongoing endogenous immune responses rather than simply filling a lymphopenic niche. Several mechanisms may explain these differences. Unlike SFB, which remains largely restricted to the intestinal epithelium (*34*), *E. gallinarum* has been detected in extraintestinal tissues of lupus-prone mice and patients (*4, 5*), suggesting broader opportunities for antigen exposure, T cell activation, and immune remodeling (*4, 9, 10, 35*). Together, these findings indicate that the microbial context, rather than broad reactivity to the microbiota, is a critical determinant of whether microbiota-reactive CD4⁺ T cells acquire pathogenic potential during autoimmunity.

A limitation of this study is the lack of tools to directly identify and track pathobiont-specific CD4⁺ T cells. While SFB-specific tetramers enabled precise analysis of commensal-reactive T cells, the contribution of pathobiont-reactive populations was inferred through microbial conditioning, functional responses, and adoptive transfer experiments rather than direct antigen-specific detection. In addition, although perturbation of cell trafficking pathways and blockade of CCR5 signaling reduced microbiota-reactive T cell accumulation and disease-associated immune responses, these approaches are not restricted to intestinal dendritic cell migration and likely affect multiple immune populations and trafficking routes. Therefore, the precise cellular and molecular pathways that drive microbiota-reactive T cell expansion during autoimmunity remain to be fully defined. Finally, the mechanisms by which pathobionts such as *E. gallinarum* promote pathogenic CD4⁺ T cell programs remain incompletely understood and may involve a combination of antigen dissemination, microbial translocation, metabolite production, and tissue-specific inflammatory cues. Addressing these questions will provide important insight into the mechanisms by which systemic autoimmunity reshapes microbiota-reactive CD4⁺ T cell immunity and may reveal opportunities for selectively targeting pathogenic microbe-specific immune responses.

Together, our findings support a model in which systemic autoimmunity broadens microbiota-reactive CD4⁺ T cell immunity through increased intestinal dendritic cell trafficking and thymic generation, while microbial context determines the functional consequences of these responses. In this framework, commensal-reactive T cells can expand and acquire effector phenotypes without substantially worsening disease, whereas the lupus-associated pathobiont *E. gallinarum* promotes effector programs that amplify autoimmune pathology. These observations suggest that microbiota-reactive T cells represent a dynamic interface between the intestinal microbiota and systemic immune responses, capable of integrating microbial and inflammatory cues. More broadly, our findings indicate that disease-associated alterations in the microbiota may influence autoimmunity not only through microbial products or translocation, but also by shaping the generation, accumulation, differentiation, and function of microbiota-reactive CD4⁺ T cells.

## MATERIALS AND METHODS

### Study Design

This study investigated how systemic autoimmunity alters microbiota-reactive CD4⁺ T cell responses and whether pathobiont-reactive T cells contribute to autoimmune disease progression. Inducible (imiquimod-treated) and spontaneous (TLR7.1 Tg) murine models of lupus-like disease were used. Mice were colonized with segmented filamentous bacteria or *Enterococcus gallinarum* and analyzed by flow cytometry, ELISA, histology, quantitative PCR, adoptive transfer, photoconversion-based cell tracking, cytokine blockade, and RNA sequencing. Primary study objectives and experimental endpoints were defined before initiation of experiments.

No formal sample size calculations were performed. Sample sizes were determined on the basis of prior experience with the experimental models and technical feasibility. Mice were randomly assigned to experimental groups to distribute age- and sex-matched animals within experiments, and littermate controls were used whenever possible. Experimental endpoints were predefined according to study design, and animals exhibiting severe clinical symptoms were euthanized in accordance with approved animal care protocols. Ex vivo restimulation and RNA sequencing experiments were performed once, whereas all other experiments were independently repeated at least twice. Investigators were blinded to experimental group allocation only during histological scoring. Outliers were identified using the ROUT method (Q = 1%) and excluded prior to statistical analysis when present. Additional details regarding biological replicates, statistical analyses, and sample sizes are provided in the corresponding figure legends.

### Mice

C57BL/6J (B6, JAX 000664), C57BL/6J-*Ptprc^em6Lutzy^*/J(*36*) (CD45.1, JAX 033076), B6.129S4-*Ifng^tm3.1Lky^*/J(*37*) (GREAT, JAX 017581), Tg(CAG-KikGR)33Hadj/J(*28*) (KikGR, JAX 013753), B6.Cg-*Rag2^tm1.1Cgn^*/J(*38*) (RAG2KO, JAX 008449), FVB-Tg(Rag2-EGFP)1Mnz/J(*39*) (RAG2-GFP, JAX 005688), and TLR7.1 transgenic (TLR7.1 Tg) lupus-prone mice were used in this study. TLR7.1 Tg mice were kindly provided by Jessica Hamerman (Benaroya Research Institute)(*17*), and RAG2-GFP mice were provided by Gretchen Diehl (Memorial Sloan Kettering Cancer Center). All other strains were originally obtained from The Jackson Laboratory and subsequently bred in-house under specific pathogen-free (SPF) conditions at the University of Virginia vivarium. All mouse strains were maintained on a C57BL/6J genetic background. Mice were used between 3 and 15 weeks of age unless otherwise indicated. Experiments included both male and female mice, and age- and sex-matched animals were used whenever possible. Littermate controls were used when available, and animals were randomly assigned to experimental groups. Mice were provided standard laboratory chow and water ad libitum and housed on hardwood chip bedding in a temperature-controlled, air-conditioned facility under a 12-hour light/dark cycle. All procedures were conducted in accordance with protocols approved by the University of Virginia Institutional Animal Care and Use Committee.

### Lupus models

#### Inducible lupus model

Mice received 25 mg of 5% imiquimod (IMQ) cream (Perrigo) applied topically to alternating left and right ears three times weekly for 6 weeks, inducing a lupus-like disease as previously described. This protocol was adapted from Zegarra-Ruiz et al., 2019 (*9, 13*)

#### Spontaneous lupus model

TLR7.1 Tg mice were used as a spontaneous model of lupus-like disease. These mice exhibit an approximately 8- to 16-fold increase in TLR7 expression, resulting in spontaneous systemic autoimmunity characterized by splenomegaly, autoantibody production, and inflammatory pathology, including glomerulonephritis, beginning at approximately 6 weeks of age (*9, 17*)

### Bacterial stock preparation and storage

Segmented filamentous bacteria (SFB) stocks were prepared from cecal contents collected under anaerobic conditions and resuspended in sterile phosphate-buffered saline (PBS) supplemented with glycerol (50% v/v). Suspensions were aliquoted and cryopreserved at −80°C.

*Enterococcus gallinarum* was cultured anaerobically at 37°C in pre-reduced Brain Heart Infusion (BHI) (Millipore Sigma) medium and expanded to the exponential phase (∼2 × 10^9^ CFU). Cultures were pooled, mixed 1:1 with BHI (60%) supplemented with glycerol (40%) and L-cysteine (0.1%) (Millipore Sigma), aliquoted, and cryopreserved at −80°C.

### Bacterial colonization

Mice were colonized with SFB using frozen stocks described above. To colonize mice with *Enterococcus gallinarum*, frozen stocks were cultured for 5 hours or until 2 × 10^9^ CFU were obtained. Cultures were normalized to 2 × 10^9^ CFU when necessary, transported on ice, and administered by oral gavage. SFB colonization was performed by two consecutive oral gavages (200 µL per mouse) on the same day, with a 6-hour interval between doses. *E. gallinarum* colonization was achieved by weekly oral gavage (200 µL per mouse).

Colonization by SFB was verified by DNA isolation and quantitative PCR analysis. Genomic DNA was isolated using a bead-beating/phenol–chloroform extraction method. Samples were lysed in DNA extraction buffer containing 200 mM Tris-HCl (pH 8.0) (Millipore Sigma), 200 mM NaCl (Promega), and 20 mM EDTA (Thermo Fisher Scientific), supplemented with 20% SDS (Thermo Fisher Scientific), and disrupted mechanically with 0.1 mm zirconia-silica beads (BioSpec Products) using a bead beater (5 × 60 s, maximum speed). Lysates were extracted with phenol:chloroform:isoamyl alcohol (Thermo Fisher Scientific) and centrifuged to separate phases. The aqueous phase was subjected to repeated organic extractions until no visible protein interface remained. DNA was precipitated with 118mM sodium acetate (Fisher Scientific) and absolute ethanol (Thermo Fisher Scientific), incubated at −80 °C, and pelleted by high-speed centrifugation (16,000 g for 5 min at 4 °C). Pellets were washed with 70% ethanol, air-dried, and resuspended in ultrapure water. Quantitative PCR was performed using SFB primers with Il23r (tissues) or 16S (fecal pellets) as housekeeping genes (Il23r F: GCAAATGGAAATGTCAGCAGAGCC; Il23r R: GCAGCTCACTTTCAGTAATCTGGG; 16S F: CGGTGAATACGTYCGG; 16S R: GGWTACCTTGTTACGACTT; SFB F: GACGCTGAGGCATGAGAGCAT; SFB R: GACGGCACGGATTGTTATTCA) (*16*).

*E. gallinarum* colonization was confirmed by culture from fecal pellets. Fecal material was homogenized through a 100 µm strainer, resuspended in sterile PBS, and inoculated into BHI medium supplemented with vancomycin (6 µg/mL) (Millipore Sigma), including pure culture and uninoculated controls. Cultures were incubated anaerobically at 37°C for 72 hours, pelleted, washed twice with sterile PBS, and resuspended prior to DNA isolation. For DNA isolation, bacterial cells were pelleted, washed in TE buffer (10 mM Tris-HCl pH 8.0, 1 mM EDTA), and resuspended. Cells were treated with lysozyme (2.5 mg/mL) (Millipore Sigma), 20% SDS, and proteinase K (0.9 mg/mL) (Millipore Sigma), followed by phenol–chloroform extraction as described above. DNA was precipitated with ethanol and sodium acetate, washed, air-dried, and resuspended in nuclease-free water (Thermo Fisher Scientific). *E. gallinarum* abundance was assessed by conventional PCR using species-specific primers (F: TTACTTGCTGATTTTGATTCG; R: TGAATTCTTCTTTGAAATCAG).

### Tissue processing and cell isolation

#### Cell isolation

Single-cell suspensions were prepared from spleen, thymus, kidney, Peyer’s patches, mesenteric lymph node (MLN), and small intestinal lamina propria. Following euthanasia, the abdominal cavity was opened, and tissues were carefully collected. Tissues other than the small intestine were mechanically dissociated through 70-µm cell strainers using a sterile plunger in complete RPMI 1640 medium (RPMI supplemented with 10% FBS) (Gibco). Cell suspensions were filtered through 40-µm strainers, centrifuged at 1,800 rpm for 5 min at 4°C, and resuspended in MACS buffer (PBS containing 2% FBS (Gibco) and 2 mM EDTA (Thermo Fisher Scientific)) unless otherwise indicated. For spleen and kidney, red blood cells were lysed using RBC lysis buffer (BioLegend) for 5 minutes at room temperature. Kidney cell suspensions were subsequently enriched by 40%/80% Percoll density gradient centrifugation (Cytiva) at 2,500 rpm for 20 min at room temperature without brake. Cells from the interphase were collected, washed, and resuspended in MACS buffer. For antigen-presenting cell (APC) and dendritic cell (DC) analysis, tissues were incubated in digestion buffer (complete RPMI supplemented with 0.01% v/v of 150 mg/mL DNase I from bovine pancreas (Millipore Sigma) and 0.01% v/v of 100 kU/mL Type 8 Collagenase from *Clostridium histolyticum* (Millipore Sigma)) and digested for 30 min at 37°C following mechanical disruption. Cell suspensions were filtered through 70-µm strainers and processed as described above.

To isolate cells from the small intestinal lamina propria, intestines were harvested and placed in ice-cold PBS. Adipose tissue and Peyer’s patches were removed, intestines were opened longitudinally, and luminal contents were discarded. Tissues were cut into approximately 2-cm segments and washed in PBS. Epithelial cells and mucus were removed by two sequential incubations at 37°C with shaking (200 rpm): first in PBS containing 1 mM dithiothreitol (DTT) (Thermo Fisher Scientific), 30 mM ethylenediaminetetraacetic acid (EDTA) (Thermo Fisher Scientific), and 10 mM HEPES for 10 min, followed by a second incubation in PBS containing 30 mM EDTA and 10 mM HEPES (Fisher Scientific) for 10 min. Tissues were then digested in Type 8 collagenase/DNase I-containing digestion buffer in complete RPMI for 1 h at 37°C with shaking. Following digestion, cells were collected and enriched by 40%/80% Percoll density gradient centrifugation as described for kidney samples. Cells were washed and resuspended in MACS buffer for downstream applications.

#### CD4^+^ T cell enrichment for tetramer-specific T cell detection

CD8⁺ T cells and CD19⁺ B cells were depleted from splenic preparations, whereas CD8⁺ T cells were depleted from thymic preparations, using magnetic-activated cell sorting. Cells were resuspended in MACS buffer at 1 × 10^8^ cells/mL and incubated with biotin-conjugated anti-CD8a (1:200, BioLegend, cat #: 100704) and anti-CD19 antibodies (1:200, BioLegend, cat #: 115504) for 15 min at 4°C. After washing, streptavidin-conjugated magnetic beads (BioLegend) were added at 5 µL per 1 × 10^7^ cells and incubated for 10 min at 4°C. Cells were then placed on a magnetic separator, and the unlabeled fraction was collected for downstream applications.

#### APC enrichment

For APC enrichment, CD90.2⁺ T cells were depleted using biotin-conjugated anti-CD90.2 antibodies and the magnetic-activated cell sorting protocol described above.

### *Ex vivo* restimulation for Nur77 analysis

#### Preparation of fecal bacterial lysate and heat-killed E. gallinarum

Fecal pellets were collected from control and IMQ-treated mice after 5 weeks of IMQ treatment and homogenized in sterile PBS containing 0.1-mm zirconia-silica beads (BioSpec Products) using a bead beater (5 × 60 s at maximum speed). The resulting suspensions were centrifuged at 2,000 rpm to remove large particles, and the supernatants were subsequently centrifuged at 10,000 × g for 10 min. The resulting pellets were resuspended in sterile PBS and heat-inactivated at 75°C for 30 min. Protein concentrations were determined by NanoDrop spectrophotometry and used to normalize lysate concentrations prior to ex vivo stimulation. For restimulation with fecal microbiota lysate, cells were incubated with 50 µg/mL lysate prepared from fecal pellets collected from mice of the same experimental group (control or IMQ-treated).

*E. gallinarum* was cultured in BHI medium for 8 h and pelleted at 3,000 × g for 10 min. Bacterial pellets were washed twice with sterile PBS and resuspended in sterile PBS to an OD_600_ of 1.0. Bacteria were then heat-killed at 75°C for 20 min.

Heat-killed *E. gallinarum* and fecal microbiota lysate were plated on BHI agar plates overnight to confirm the absence of bacterial growth.

#### *Ex vivo* T cell restimulation with fecal bacterial lysate, SFB peptide or *E. gallinarum*

After single-cell suspensions were prepared from the spleen or MLN, splenocytes were seeded in 48-well plates at 5 × 10^6^ cells/well, and MLN cells were seeded in 96-well plates at 1 × 10^6^ cells/well. For restimulation with fecal bacterial lysate, cells were incubated with 50 µg/mL fecal bacterial lysate prepared from the same mouse (autologous stimulation) for 4 h. For restimulation with SFB peptide or heat-killed *E. gallinarum*, cells were incubated with 1 µg/mL SFB3340 peptide (QFSGAVPNKTD (*15*); GenScript) or heat-killed *E. gallinarum* at a multiplicity of infection (MOI) of 50, respectively, for 4 h. Cells were subsequently processed for flow cytometric analysis of Nur77 expression.

### Flow cytometry and tetramer staining

Cells isolated from spleen, kidneys, thymus, MLN, Peyer’s patches, or small intestinal lamina propria were surface-stained in MACS buffer or Fc/Block buffer for 25 minutes in the dark at 4 °C with fluorescently conjugated antibodies.

#### Tetramer staining

Cells were stained with PE- and APC-conjugated tetramers diluted in Fc/Block buffer (1:300 anti-mouse CD16/CD32 in MACS buffer) to a final concentration of 10nM. Tetramers were used to detect antigen-specific CD4^+^ T cells: SFB, SFB3340 (*15*) 200-210: QFSGAVPNKTD; bacillus flagellin 464–472: YSNANILSQ; and Human DRB1*08:03 protein CLIP 87-101: PVSKMRMATPLLMQA (NIH Tetramer Core Facility). Cells were incubated at room temperature for 20 min, followed by 40 min incubation at 4 °C in the dark. Next, surface antibody staining was performed.

#### T cell phenotyping

For T cells, we used antibodies specific to CD3 (1:200, BioLegend, cat #: 100229), CD4 (1:300, BioLegend, cat #: 100548), CD8a (1:300, BioLegend, cat #: 100752), TCRβ (1:200, BioLegend, cat #: 109241), CD44 (1:200, BioLegend, cat #: 103028), CD62L (1:200, BioLegend, cat #: 104445), CXCR5 (1:200, BioLegend, cat #: 145526), Ki67 (1:100, BioLegend, cat #: 652410), PD1 (1:200, BioLegend, cat #: 135231), and CD45 (1:400, BioLegend, cat #: 103149). For identification of T cell subsets, after staining for surface antigens, cells were fixed and permeabilized using an eBioscience^TM^ Foxp3/ Transcription Factor Fixation/Permeabilization Concentrate and Diluent kit (Thermo Fisher Scientific) overnight at 4 °C and stained with antibodies against FOXP3 (1:100, Thermo Fisher Scientific, cat #: 56-57773-82), RORγt, T-bet (1:100, BioLegend, cat #: 644828), Bcl-6 (1:100, BioLegend, cat #: 358514), and Nur77 (1:100, Fisher Scientific, cat #: 50-110-28 and 50-112-8996) for 1 hour at room temperature.

#### APC and DC phenotyping

For APC phenotyping, antibodies against CD11b (1:400, BioLegend, cat #: 101226), CD11c (1:400, BioLegend, cat #: 117328), Ly6G (1:400, BioLegend, cat #: 127614), I-A/I-E (1:400, BioLegend, cat #: 107641), CD19 (1:300, BioLegend, cat #: 115521), B220 (1:300, BioLegend, cat #: 103244), CD45 (1:400, BioLegend, cat #: 103149), CD103 (1:200, BioLegend, cat #: 121406), CX3CR1 (1:200, BioLegend, cat #: 149027), Ly6C (1:200, BioLegend, cat #: 128024) and PDCA-1 (1:300, BioLegend, cat #: 127039) were used. For DC phenotyping, antibodies against F4/80 (1:300, BioLegend, cat #: 123133), CD64 (1:200, BioLegend, cat #: 139309), CD88 (1:200, BioLegend, cat #: 135809), CCR7 (1:200, BioLegend, cat #: 120106), XCR1 (1:200, BioLegend, cat #: 148218), and SIRPα (1:200, BioLegend, cat #: 144016) were used. For plasma cells, antibodies against CD138 (1:200, BioLegend, cat #: 142521) and IgG (1:400, Thermo Fisher Scientific, cat #: A16067) were used. In all cases, viability was assessed using a Zombie Violet™ Fixable Viability Kit (1:300, BioLegend, cat #: 423114).

#### Gating strategy: T cells

After exclusion of dead cells and doublets, tetramer-specific CD4^+^ T cells were defined as CD45^+^ TCRβ^+^ CD3^+^ CD4^+^ CD8^-^ and PE- and APC-Tetramer^+^. Th1, Th17, regulatory T cell (Treg), and RORγt^+^ Treg populations were identified as T-bet^+^ (Th1), RORγt^+^ (Th17), FOXP3^+^ (Tregs), and RORγt^+^ FOXP3^+^ (RORγt^+^ Tregs) after gating out nonviable cells and doublets and selecting for CD4^+^ T cells. Following viability and singlet discrimination, Tfh cells were defined as BCL6⁺ CXCR5⁺ PD-1⁺ FOXP3⁻ CD4^+^ T cells, whereas Tfr cells were identified using the same gating strategy, except for FOXP3 positivity. Cytokine-producing populations were defined by expression of TNFα (1:100, BioLegend, cat #: 506346), IL-21 (1:100, Fisher Scientific, cat #: 50-111-72), IL-17A (1:100, BioLegend, cat #: 506927), or IFNγ (1:100, BioLegend, cat #: 505836) within CD4^+^ T cells.

#### Gating strategy: APCs and DCs

After exclusion of dead cells and doublets, B cells were identified as CD45^+^ CD19⁺ B220⁺ MHCII⁺ cells, plasma cells were defined as CD45^+^ CD19⁻ B220^int^ MHCII⁺ CD138^+^ cells, and plasmacytoid dendritic cells (pDCs) as CD45^+^ CD19⁻ B220⁺ CD11b⁻ PDCA1⁺ CD11c⁺ cells. DCs were defined as CD45^+^ B220⁻ CD19⁻ CD11b⁺ Ly6C^-^ CD11c⁺ MHCII⁺ cells and subdivided into SIRPα⁺ and XCR1⁺ subsets. Additional APC populations were defined as CD11c⁺MHCII⁻CX3CR1⁺ (transitional monocytes), CD11c⁻MHCII⁻CX3CR1⁺ (classical monocytes), CD11c⁻MHCII⁺CD64⁺CX3CR1⁺ (macrophages), and CD11c⁺MHCII⁺ (monocyte-derived dendritic cells), after CD45^+^ CD19⁻ B220⁻ CD11b⁺Ly6C⁺ selection.

Data were acquired using a Cytek Aurora Full Spectral 5-Laser flow cytometer (Cytek Biosciences) or an Attune NxT flow cytometer (Thermo Fisher Scientific) and analyzed using FlowJo v10 (Tree Star). Absolute cell numbers were determined using Precision Count Beads (BioLegend).

### In vitro cytokine T cell stimulation

Single-cell suspensions were prepared, and 1 × 10^6^ cells per well were added to 48-well plates. Cells were stimulated with a phorbol 12-myristate 13-acetate (PMA; 50 ng/mL) (Millipore Sigma) and ionomycin (1000 ng/mL) (Millipore Sigma) in the presence of Brefeldin A (5μg/mL) (BioLegend) for 4 hours before surface and tetramer staining. Fixation, permeabilization, and intracellular cytokine staining were performed using the eBioscience^TM^ Foxp3/ Transcription Factor Fixation/Permeabilization Concentrate and Diluent (Thermo Fisher Scientific).

### Transcriptome data processing and analysis

#### T cell isolation and RNA extraction

Approximately 10,000–20,000 T cells were fluorescence-activated cell sorting (FACS)-sorted from the thymus or spleen. Both SFB tetramer-negative and tetramer^+^ T cells were sorted directly into TRIzol LS (Thermo Fisher Scientific) at room temperature and immediately flash-frozen. SFB tetramer^+^ T cells were identified as viable singlet populations of MHCII⁻, CD19⁻, CD8⁻, CD3^+^, and CD4^+^ cells and isolated using peptide–MHC class II tetramers, enabling the selective purification of antigen-specific T cell populations. For each biological replicate, cells from four mice were pooled prior to downstream processing. Total RNA isolation, RNA-seq library preparation, and sequencing were performed by Azenta Life Science.

#### RNA-seq preprocessing and exploratory analysis

Raw paired-end reads were processed using BBDuk (BBMap suite v38.84) (*40*) to remove adapter sequences and low-complexity reads. Adapter trimming was performed using standard reference sequences with a k-mer length of 23 and a minimum k-mer length of 11 (k=23, mink=11, hdist=1). Paired-end overlap trimming (tbo) and trimming based on pair length (tpe) were enabled to enhance accuracy. Subsequently, reads were filtered for low complexity using an entropy threshold of 0.5 (entropy=0.5). Transcript quantification was performed from cleaned reads using. Salmon (v1.10.3) (*41*) in mapping-based mode. The reference transcriptome index was constructed from GENCODE (*42*) mouse annotation release vM37 using the GRCm39 primary genome assembly. The full genome sequence was included as decoy sequences to enable selective alignment. GC content bias correction was enabled using the --gcBias option. Transcript-level abundance estimates were aggregated to gene-level counts using the tximport R package (*43*) Gene-to-transcript mappings were derived from the GENCODE vM37 primary assembly annotation file, retaining entries annotated as transcripts.

Gene-level count matrices were imported into DESeq2 using DESeqDataSetFromTximport. To focus on biologically relevant transcripts, analyses were restricted to genes annotated as protein-coding or immune receptor genes, including T cell receptor (TR) and immunoglobulin (IG) gene families (protein_coding, TR_V, TR_J, TR_D, TR_C, IG_V, IG_LV, IG_J, IG_D, IG_C). Genes with low expression were filtered based on mean raw counts, using thresholds of greater than 10. Size factor normalization was performed using DESeq2 prior to downstream analyses. For dimensionality reduction and exploratory analysis, variance-stabilizing transformation (VST) was applied to normalized count data. Gene-wise variances were computed, and the top 1,000 most variable genes were selected. Principal component analysis (PCA) was performed using the prcomp function on the transposed expression matrix. Percent variance explained by each principal component was calculated and used for axis labeling in PCA visualizations.

#### Differential gene expression analysis and functional enrichment analyses

Differential gene expression analyses were conducted using DESeq2 (v1.42.1) (*44*) For each comparison, samples were subset to include only the relevant experimental groups. The DESeq2 model was fitted using standard settings. Group reference levels were set explicitly for each comparison. Differential expression results were extracted using Wald tests with Benjamini–Hochberg correction for multiple hypothesis testing. Genes were considered significantly differentially expressed based on an adjusted P value ≤ 0.05. Z-score-normalized gene counts were used for reporting and visualization in volcano and heatmap plots.

#### Curated immune-program signature analysis

To assess coordinated transcriptional programs, we assembled curated gene sets using mouse gene symbols for seven T cell programs from canonical lineage and functional markers, plus a type I interferon signature. Th1 (Tbx21, Ifng, Stat1, Stat4, Cxcr3, Eomes, Nkg7, Gzmb, Gzmk, Bhlhe40, Ccl5), Tfh (Bcl6, Cxcr5, Pdcd1, Icos, Il21, Ascl2, Tox2, Tox, Maf, Ikzf2, Sh2d1a), TCR signaling (Cd3e, Zap70, Lck, Malt1, Itpkb, Nr4a1, Nr4a2, Egr2, Fos, Junb, Dusp1, Dusp4, Jak2, Rasa1), T cell activation (Cd69, Il2ra, Tnfrsf9, Tnfrsf4, Cd28, Ctla4, Lag3, Tigit, Tnfsf8, Nfkbiz, Klf6, Ccl3, Ccl4), Proliferation (Mki67, Top2a, Ccnb1, Cdk1, Cdk6, Cdc6, Cdc45, Aurkb, Plk1, Rrm2, Birc5, Ezh2, Nusap1), Cytokine effector (Il2, Il10, Tnf, Lta, Ltb, Tgfb1, Cxcl10, Il4), Naïve quiescence (Sell, Ccr7, Lef1, Tcf7, Klf2, S1pr1, Bach2, Foxp1, Dntt, Il4ra, Selplg), Type I IFN (Irf7, Isg15, Oas1a, Oasl2, Oas2, Oas3, Stat1, Stat2, Ifit1, Ifit3, Mx1, Mx2, Rsad2, Usp18, Ifi44, Isg20, Bst2, Irf9, Ifih1, Xaf1, Cmpk2, Ifi204, Ifi211). Each gene was assigned to a single program to keep the sets mutually exclusive.

#### Signature-level Gene Set Enrichment Analysis

GSEA was performed with the fgsea package (v1.24.0; R v4.2.1). For each contrast, genes were ranked by the DESeq2 Wald statistic from the differential-expression table computed on all detectably expressed genes. Duplicate gene symbols were collapsed to the entry with the largest absolute statistic, and ranks were sorted in decreasing order. Enrichment was computed against the curated signatures using fgsea with minSize = 3, maxSize = 500, and eps = 0. Significance was assessed using Benjamini–Hochberg-adjusted P values. Normalized enrichment scores were displayed as bar plots, with bar length and fill representing the normalized enrichment score and the adjusted P value annotated for each bar.

Marker-gene bar for Ifng, Il21, Maf, and Bhlhe40 show DESeq2 size-factor-normalized counts per group with individual replicate values overlaid. Significance reflects the DESeq2 adjusted P value for that contrast.

### RNA extraction and quantitative real-time PCR

Spleen sections were collected in TRIzol (Thermo Fisher Scientific) containing 1.0 mm zirconium-silicate beads (BioSpec Products) and either stored at −80°C or processed immediately. Samples were lysed by bead beating (3 × 60 s, maximum speed) with cooling on ice between cycles. RNA was extracted by chloroform (Millipore Sigma) phase separation, and the aqueous phase was precipitated with isopropanol (Millipore Sigma), washed twice with 75% ethanol, briefly air-dried, and resuspended in ultrapure water, according to the manufacturer’s protocol. RNA concentration and purity were assessed using a NanoDrop. cDNA synthesis was performed using iScript™ cDNA Synthesis Kit (Bio-Rad), and gene expression was quantified by qPCR using iTaq Universal SYBR Green (Bio-Rad) and gene-specific primers for Irf7, Cxcl9, Oas3, Oas1a, Stat1, and Isg15, with HPRT as the housekeeping gene (Irf7 F: ACAGCACAGGGCGTTTTATC; Irf7 R: GAGCCCAGCATTTTCT CTTG; Cxcl9 F: GGAGTTCGAGGAACCCTAGTG; Cxcl9 R: GGGATTTGTAGTGGATCGTGC; Oas3 F: CGGAGCTCTGGATAAGCTGG; Oas3 R: GGTGATGTTTAGCGACCCCA; Oas1a F: ATCAGCCGTCAATGTCGTGT; Oas1a R: CTTCACCACCTTGGAGACCC; Stat1 F: TCACAGTGGTTCGAGCTTCAG; Stat1 R: GCAAACGAGACATCATAGGCA; Isg15 F: GGTGTCCGTGACTAACTCCAT; Isg15 R: CTCCAGCCAGAACTGGTCTTC; HPRT F: CAAACTTTGCTTTCCCTGGT; HPRT R: TCTGGCCTGTATCCAACACTTC). Reactions were run on a QuantStudio 6 Flex (Applied Biosystems). Gene expression was normalized to HPRT and is presented as fold change relative to the indicated control group.

### Enzyme-linked immunosorbent assays (ELISAs)

Total IgG and autoantibodies against double-stranded DNA (dsDNA) and single-stranded RNA (ssRNA) in serum were quantified by ELISA assay.

#### Total IgG ELISA

96-well plates were coated overnight at room temperature in the dark with goat anti–mouse IgG (SouthernBiotech; 0.1 µg/mL) diluted in ELISA coating buffer (Bio-Rad). Plates were washed with PBS containing 0.05% Tween-20 (Millipore Sigma) and blocked with coating buffer supplemented with 2% Bovine Serum Albumin (BSA) (Tocris Bioscience) for at least 90 min at room temperature. After additional washes, serum (1:10,000) was added in dilution buffer (PBS containing 1% BSA). A standard curve was generated using unconjugated mouse IgG (SouthernBiotech) serially diluted 1:2 in dilution buffer, starting at 50 ng/mL. Plates were incubated for 90 min at room temperature, washed, and incubated with HRP-conjugated goat anti–mouse IgG secondary antibody (Southern Biotech; 1:10,000) for 90 min at room temperature. Following extensive washing, plates were developed using TMB substrate (Neogen) at room temperature in the dark, and the reaction was stopped with 2M sulfuric acid (Fisher Scientific).

#### Anti-dsDNA ELISA

Plates (High Binding, Corning) were pre-coated with protamine sulfate (100 µL per well at 150 µg/mL) and incubated for 45 minutes at room temperature in the dark. Wells were rinsed twice with ultrapure water, blotted, and air-dried. Plates were then coated with 100 µg/mL salmon sperm DNA (Thermo Fisher Scientific) and blocked with PBS containing 3% BSA and 5% fetal bovine serum (FBS). Serum samples were diluted 1:1000 in dilution buffer and incubated for 1 hour at 37°C. Wells were washed and incubated with rabbit anti–mouse IgG conjugated to HRP (0.1 µg/mL, Fisher Scientific), for 1 hour at 37°C. Color development was performed using TMB substrate, and the reaction was stopped with 2M sulfuric acid.

#### Anti-ssRNA ELISA

Plates were pre-coated with protamine sulfate (100 µL per well at 50 µg/mL) and processed as above. Plates were coated with 5 mg/mL yeast RNA (Thermo Fisher Scientific) and blocked with PBS containing 0.05% Tween-20 and 5% FBS. Serum samples were diluted 1:1000 in dilution buffer and incubated for 1 hour at 37°C. Subsequent washing, secondary antibody incubation, and development steps were performed as described for the anti-dsDNA ELISA.

Absorbance was measured at 450 nm with a 570 nm reference wavelength.

### Histology

Kidneys were harvested from mice across the indicated experimental conditions. All samples were fixed in 10% neutral buffered formalin, transversely sectioned, and stained with hematoxylin and eosin (H&E). Renal histopathology was evaluated using a modified scoring system based on the Inotiv mouse lupus nephritis protocol, originally described by Chan, Madaio, and Shlomchik and subsequently adapted by Bendele (*45*). Five histopathological parameters were assessed. Glomerular involvement was determined by measuring the diameters of 20 representative glomeruli per kidney and calculating the mean value, with scores ranging from 0 (normal; ≤65 µm) to 5 (severe diffuse proliferative; >110 µm). Crescent formation was scored according to the proportion of glomeruli exhibiting crescents, ranging from 0 (absent) to 5 (>50% of glomeruli affected). Protein cast accumulation in cortical tubules was graded based on the percentage of cortex involved, from 0 (normal) to 5 (severe; >75%). Interstitial nephritis, defined as inflammatory infiltrates independent of vascular structures, was scored for the extent and distribution of mononuclear cell infiltration in the renal pelvis and cortex, from 0 (normal) to 5 (diffuse involvement, 76–100%). Vasculitis was assessed by evaluating the number and severity of perivascular inflammatory lesions and the presence of vessel wall necrosis, with scores ranging from 0 (normal) to 5 (severe multifocal disease with necrosis). Individual scores for each parameter were summed to generate a composite renal pathology score. Histological evaluation was performed in a blinded manner.

### Adoptive transfer protocols

#### T and B cell isolation

Single-cell suspensions were prepared from spleen and thymus. Tissues were harvested under sterile conditions, transferred to ice-cold MACS buffer, and mechanically dissociated through a 70 µm cell strainer using a sterile syringe plunger. Cell suspensions were further filtered through a 40 µm strainer, centrifuged at 300 × g for 5 minutes at 4°C, and erythrocytes were lysed by incubation in RBC lysis buffer for 5 minutes at room temperature. Cells were washed with cold PBS and resuspended in MACS buffer. Cell numbers and viability were determined by Trypan Blue (ThermoFisher Scientific) exclusion using a hemocytometer, and cell concentrations were adjusted to 1 × 10^8^ cells/mL.

For total CD4⁺ T cell isolation, cells were incubated with a biotinylated antibody cocktail followed by streptavidin-conjugated magnetic nanobeads for negative selection, according to the manufacturer’s instructions (BioLegend). After magnetic separation, the unlabeled fraction containing CD4⁺ T cells was collected (negative selection), washed, and pooled. Cell counts and viability were assessed, and suspensions were adjusted to 5 × 10^6^ cells/mL in PBS.

For B cell isolation, cells were incubated with anti-CD19–conjugated magnetic nanobeads (BioLegend) for 15 minutes on ice. Cells were washed, resuspended in MACS buffer, and placed on a pre-chilled magnetic separator. The unlabeled fraction was discarded, and the labeled CD19⁺ cells were recovered following two rounds of magnetic separation (positive selection). Purified B cells were pooled, counted, and resuspended in PBS at a final concentration of 5 × 10^6^ cells/mL. Purified cells were maintained at 4°C and used immediately for intraperitoneal injection.

#### For transfer into RAG2KO mice

SFB-colonized mice: B6 donor mice were treated topically with 5% IMQ for 6 weeks, while control donors remained untreated. At week 4 of the treatment period, both donor groups were colonized with SFB as described before. Two weeks after colonization, donor mice were euthanized, and splenic CD4⁺ T cells were isolated as described above. RAG2KO recipient mice were colonized with SFB one week prior to cell transfer. All recipient mice received splenic B cells (5 × 10^5^ cells) isolated from untreated control donors, together with splenic CD4⁺ T cells (5 × 10^5^ cells) derived either from untreated or IMQ-treated donors. Following cell transfer, all recipients received IMQ treatment for 6 weeks before tissue collection.

*E. gallinarum-*colonized mice: Transfer experiments with pathobionts occurred as above, with the following modifications. In the donors, *E. gallinarum* was administered twice, at weeks 4 and 5 of IMQ treatment. Recipient mice were colonized with *E. gallinarum* one week prior to adoptive transfer and maintained by weekly gavage throughout the experiment.

#### For transfer into CD45.2 mice

CD45.1 mice were used as cell donors and treated topically with 5% IMQ for a total of 6 weeks. During IMQ treatment, donor mice were either maintained uncolonized or orally colonized with *Enterococcus gallinarum*, as above. Two weeks after colonization, donor mice were euthanized, and splenic CD4⁺ T cells (1 × 10^6^ cells) were transferred into CD45.2⁺ recipient mice. *E. gallinarum*–colonized recipients received cells from uncolonized or *E. gallinarum*–colonized donors. Colonization of recipient mice was maintained by weekly oral gavage. Recipient mice received IMQ treatment beginning 4 weeks before adoptive transfer, which was continued for an additional 2 weeks following transfer until tissue collection.

#### For transfer to assess cytokine blocking

For cytokine-blocking experiments, adoptive transfer studies were performed using the *E. gallinarum*–colonized transfer protocol described above. Following transfer, RAG2KO recipients were treated with IMQ for 6 weeks. Mice received intraperitoneal injections of vehicle (PBS), anti-mouse IL-21R monoclonal antibody alone, or a combination of anti-mouse IL-21R (InVivoMAb, cat#: BE0258) and anti-mouse IFNγ (InVivoMAb, cat#: BE0055) monoclonal antibodies (200 µg per mice). Antibody treatments were initiated 2 weeks after adoptive transfer and administered twice weekly until the end of the experiment.

### Pertussis toxin (PTx) administration

To assess the contribution of PTx–sensitive pathways, mice underwent a six-week course of topical 5% imiquimod treatment. Following four weeks of imiquimod exposure, animals were orally colonized with SFB. Pertussis toxin (Tocris Bioscience) was administered intraperitoneally 24 h before SFB inoculation, whereas control mice received vehicle only. PTx was reconstituted in sterile PBS containing 0.2% BSA to a final stock concentration of 1 mg/mL, aliquoted, and stored at −80 °C. Fresh working solutions were prepared on the day of use to deliver 300 ng per mouse in a total volume of 200 µL. Injections were performed on days 28 and 35 of the protocol.

### CCR5 blockade

For CCR5 blockade experiments, mice received topical 5% imiquimod for six weeks and were orally colonized with segmented filamentous bacteria (SFB) during the final two weeks of treatment. Beginning one day prior to SFB colonization, mice were administered intraperitoneal injections (100 μL) of TAK-779 (Sigma-Aldrich), a small-molecule CCR5 antagonist. TAK-779 was administered at 150 μg per dose in 5% mannitol (Sigma-Aldrich) every other day for a total of five doses. Thus, CCR5 blockade was performed throughout the final 1.5 weeks of imiquimod treatment, coinciding with SFB colonization and the period of microbiota-reactive T cell expansion.

### Photoconversion and intestinal immune cell tracking (KikGR Mice)

The cecum of Tg(CAG-KikGR)33Hadj/J (KikGR, JAX 013753) mice was photoconverted using a 405 nm violet laser (peak power, 5 mW) applied for 5 minutes. Mice were anesthetized with 2.5% isoflurane (vol/vol) in 2 L/min oxygen and maintained at 37°C for the duration of the procedure. A 3 cm midline abdominal incision was performed to exteriorize the cecum, which was continuously hydrated with sterile saline. Following photoconversion, the cecum was repositioned into the abdominal cavity, and the incision was closed with sutures. Twenty-four hours after photoconversion, tissues were harvested and analyzed by flow cytometry to detect photoconverted RFP⁺ cells.

### Statistical analysis

All statistical analyses were performed using GraphPad Prism (v10.6.1). Individual data points are shown alongside the mean. “n” represents the number of biological samples analyzed in each experiment and the value of n is indicated in the corresponding figure legend. Differences between two groups were evaluated using a two-tailed unpaired Mann–Whitney U test. For comparisons involving more than two groups, a Kruskal–Wallis test was used. Two-way ANOVA was used for experiments involving comparisons across multiple groups and populations. Multiple-comparison testing was corrected using Sidak’s method. Outliers were identified using the ROUT method (Q = 1%) and excluded prior to statistical analysis. A P value < 0.05 was considered statistically significant. Statistical significance is indicated as follows: ns, not significant; \**P* ≤ 0.05; \*\**P* ≤ 0.01; \*\*\**P* ≤ 0.001; and \*\*\*\**P* ≤ 0.0001.

## ACKNOWLEDGMENTS

We thank the NIH Tetramer Core Facility (contract number 75N93020D00005) for providing I-Ab tetramers specific for SFB3340 (200–210; QFSGAVPNKTD) and Bacillus flagellin (464–472; YSNANILSQ) and Human DRB1*08:03 protein (87-101; PVSKMRMATPLLMQA). Flow cytometry and cell sorting were performed in the University of Virginia (UVA) Flow Cytometry Core Facility (RRID: SCR_017829), supported in part by NCI grant P30-CA044579; cell sorting on the Sony MA900 CS was enabled through NIH S10 instrumentation grant 1S10OD034355-01. Histological processing was performed by the UVA Research Histology Core Facility (RRID: SCR_025470). We thank Jessica Hamerman (Benaroya Research Institute) for TLR7.1 transgenic mice, Martin Kriegel (University of Münster) for *Enterococcus gallinarum* isolates, and Gretchen Diehl (Memorial Sloan Kettering Cancer Center) for RAG-GFP mice and initial fecal pellets from SFB-monocolonized mice. We are also grateful to Gretchen Diehl and Amanda Chen (Memorial Sloan Kettering Cancer Center) for guidance on the use of KikGR mice. Finally, we thank all members of our laboratory, as well as members of the Beirne B. Carter Center for Immunology Research and the Department of Microbiology, Immunology, and Cancer Biology at UVA, for mentoring and helpful feedback throughout this study.

## Data, code, and materials availability

The mRNA-seq data generated in this study were deposited in the NCBI Gene Expression Omnibus (GEO) database under accession GSE315533. All materials are commercially available or were obtained via material transfer agreement from outside sources as described in Materials and Methods. This paper does not report any original code. All data needed to support the conclusions of the paper are present in the paper or the Supplementary Materials. Any data requests will be fulfilled by the lead contact.

## AUTHOR CONTRIBUTIONS

Conceptualization and supervision were performed by D.F.Z.-R., who also led funding acquisition, visualization, and writing of the original draft and revisions. Data acquisition was carried out by M.J.V.-A., M.K., S.R.D.-P., K.S.S.-M., J.D., and G.L.M. Data analysis was conducted by M.V.-A., M.K., S.R.D.-P., K.S.S.-M., L.F.S.-U. (RNA-seq), and S.S.J. (histopathology). Visualization and figure preparation were performed by M.J.V.-A., M.K., S.R.D.-P., G.L.M., and D.F.Z.-R. The original manuscript was written by M.V.-A., M.K., S.R.D.-P., and D.F.Z.-R., and reviewed and edited by M.K., S.R.D.-P., L.F.S.-U., G.L.M., and D.F.Z.-R.

## SUPPLEMENTARY MATERIALS

**Figure S1:**
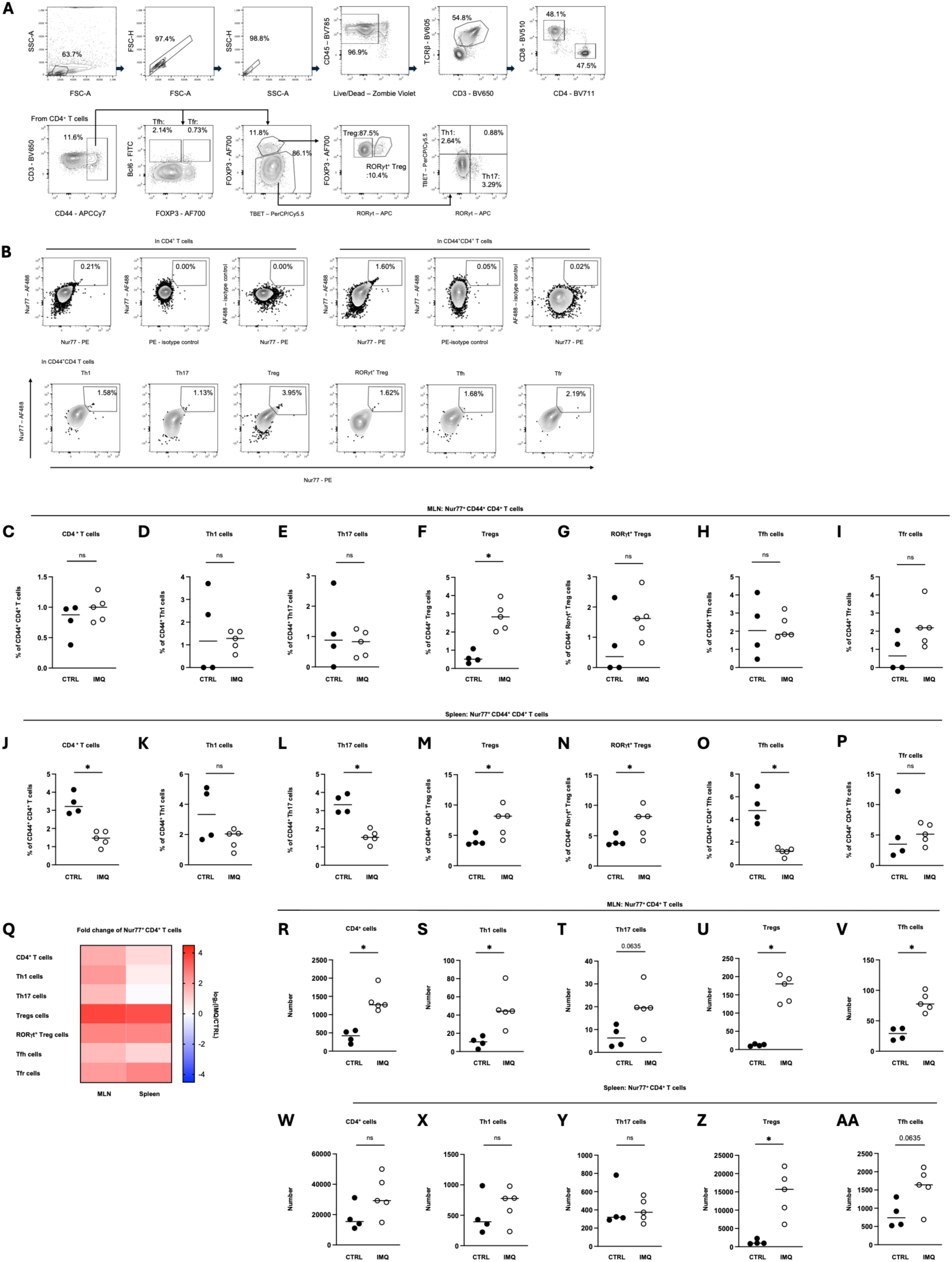
Gating strategy and extended analysis of microbiota-reactive CD4⁺ T cell responses. (A) Gating strategy for ex vivo restimulation assay. Nur77 expression was analyzed in Th1, Th17, Treg, RORγt⁺ Treg, Tfh, Tfr, and cells from the mesenteric lymph nodes (MLN) and spleen, gated from total CD4⁺ T cells or CD44⁺ CD4⁺ T cells. (B) Nur77 expression across T cell subsets and isotype control for Nur77-PE and Nur77-AF488. (C-P) Frequencies of Nur77-expressing CD44^+^ CD4^+^ (C, J), Th1 (D, K), Th17 (E, L), Treg (F, M), Rorγt^+^ Treg (G, N), Tfh (H, O), and Tfr (I, P) cells in the mesenteric lymph nodes (C-I) or spleen (J-P) of CTRL and IMQ mice (n = 4-5). (Q) Heatmap depicts log_2_(IMQ/CTRL) fold change for Nur77⁺ CD4⁺ T cells in MLN and spleen (n = 4-5). (R-AA) Counts of Nur77-expressing CD4^+^ (R, W), Th1 (S, X), Th17 (T, Y), Treg (U, Z), and Tfh (V, AA), in MLN (R-V) or spleen (W-AA) of CTRL and IMQ mice (n = 4-5). Data shown as means. Statistical significance was determined by unpaired two-tailed Mann-Whitney U test. *\*p < 0.05, **p < 0.01, ***p < 0.001, ****p < 0.0001.* See also Figure 1.

**Figure S2:**
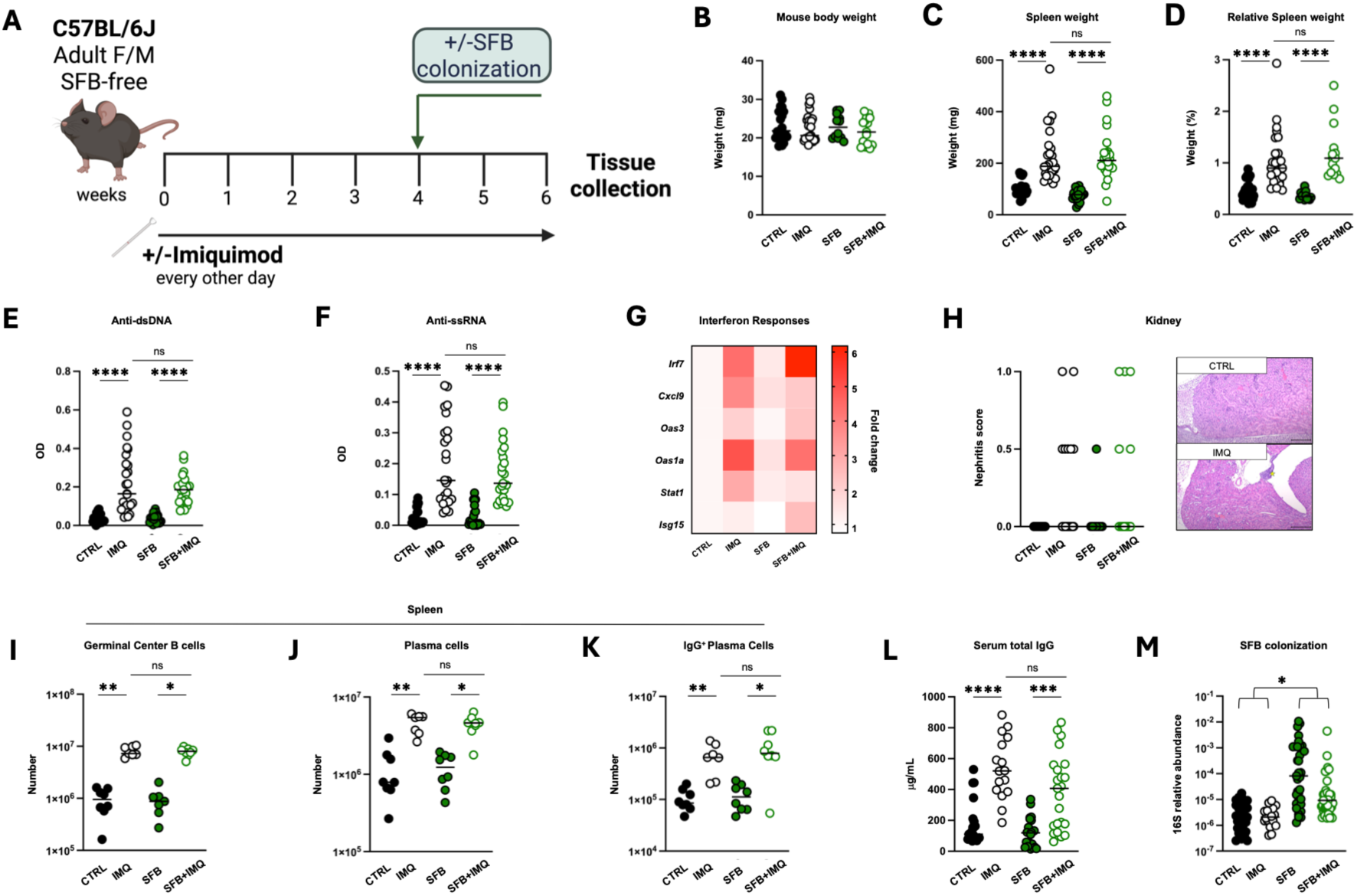
TLR7-driven autoimmune pathology develops independently of SFB colonization. (A) Experimental schematic of topical imiquimod (IMQ) administration and segmented filamentous bacteria (SFB) colonization. Mice received IMQ topically three times per week for six weeks and were colonized with SFB after four weeks of IMQ treatment. (B-D) Body weight (B), spleen weight (C), and spleen/body weight ratio (D) in CTRL, IMQ, SFB, and SFB+IMQ mice (n = 12-27). (E and F) Serum anti-dsDNA IgG (E) and anti-ssRNA IgG (F) (n = 20-25). (G) Type I IFN-related gene expression in spleen (log₂ fold change normalized to CTRL, n = 8-13). (H) Minimal interstitial mononuclear infiltrates in both groups on histopathology scoring. Hematoxylin and eosin stain (HE) (n = 10-23). (I-K) Splenic germinal center B cells (I), plasma cells (J), and IgG⁺ plasma cells (K) (n = 7-8) (L) Serum total IgG (n = 16-23). (M) qPCR of SFB tetramer^+^ CD4⁺ T cells DNA normalized to total 16S gene abundance in mouse fecal pellets (n = 18-40). Panel A was created with Biorender.com. Data shown as means. Scale bar = 0.1 mm. Statistical significance was determined by Kruskal-Wallis test (B-L) or two-way ANOVA (M). *\*p < 0.05, **p < 0.01, ***p < 0.001, ****p < 0.0001.* See also Figure 2.

**Figure S3:**
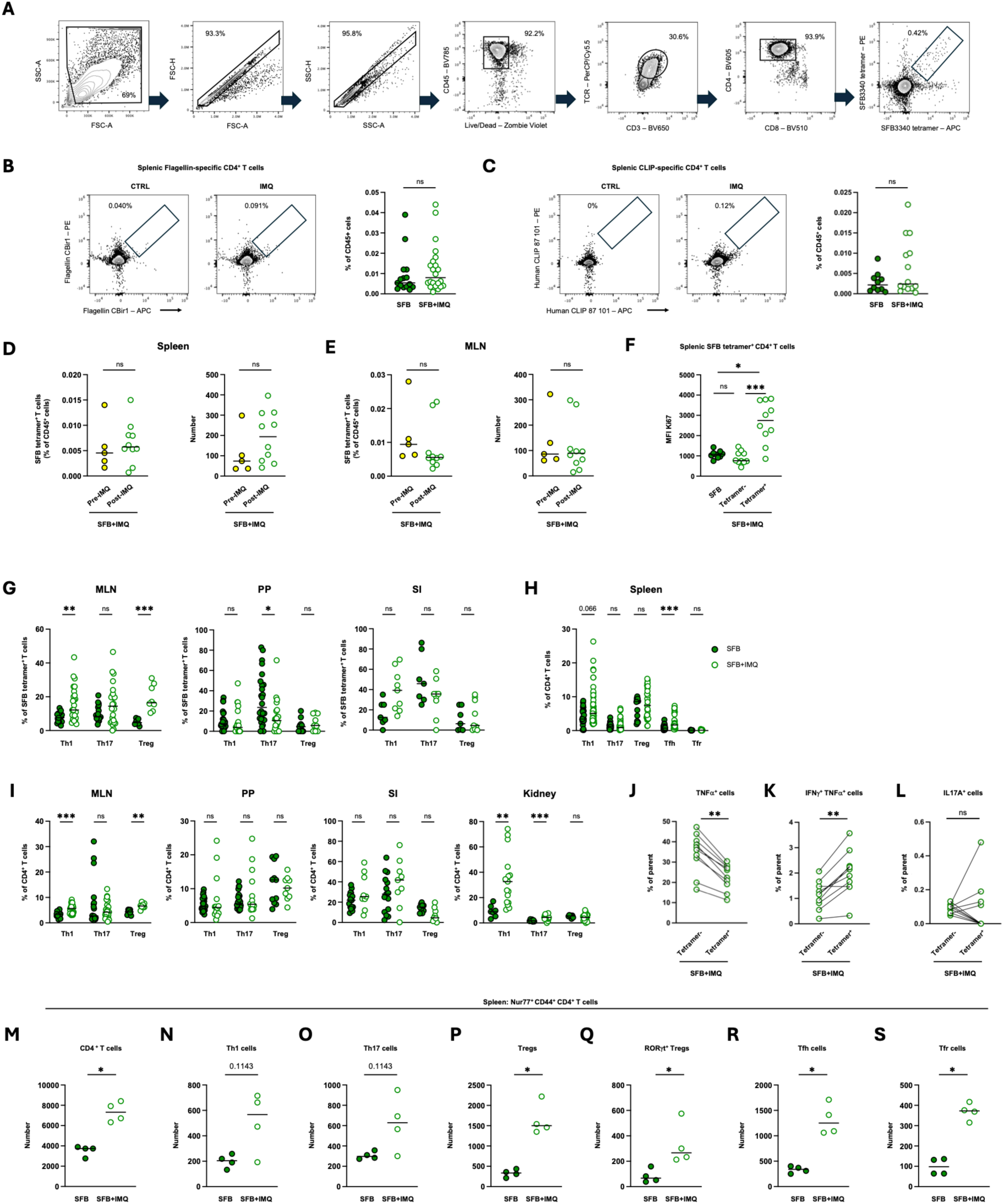
Characterization of SFB tetramer⁺ CD4⁺ T cells across tissues. (A) Gating strategy for SFB tetramer^+^ CD4⁺ T cells. (B and C) Frequencies of flagellin tetramer^+^ (B) and CLIP tetramer^+^ (C) CD4⁺ T cells in spleen of SFB-colonized (SFB) or SFB-colonized imiquimod-treated (SFB+IMQ) (n = 10-22). (D and E) SFB tetramer^+^ CD4^+^ T cells in spleen (D) and mesenteric lymph node (MLN) (E) of IMQ-treated mice with SFB colonization prior to and after IMQ treatment (n = 5-10). (F) Mean fluorescence intensity (MFI) of Ki67 in total CD4^+^ T cells from SFB and SFB tetramer^-^ and SFB tetramer^+^ CD4^+^ T cells from SFB+IMQ mice (n = 10 for all groups). (G) Effector subsets of SFB tetramer^+^ T cells in MLN, Peyer’s Patches (PP), or small intestinal lamina propria (SI) of SFB and SFB+IMQ mice (n = 8-26) (H) Effector T cell subsets in the spleen of SFB and SFB+IMQ mice (n = 14-45) (I) Effector subsets of total CD4^+^ T cells in MLN, PP, SI, or kidney of SFB or SFB+IMQ mice (n = 8-26) (J-L) TNFα^+^ (G), TNFα^+^ IFNγ⁺ (H) and IL-17A⁺ (I) production in SFB tetramer^-^ and SFB tetramer^+^ CD4⁺ T cells in the spleen of SFB+IMQ mice (n = 10 for all groups). (M-S) Nur77^+^ CD4^+^ (M), Th1 (N), Th17 (O), Tregs (P), RORγt^+^ Tregs (Q), Tfh (R), or Tfr (S) cells in spleen of SFB and SFB+IMQ mice (n = 4). Data shown as means. Statistical significance was determined by Kruskal-Wallis test (B-E, M-S), unpaired two-tailed Mann-Whitney U test (F-I), or two-tailed Wilcoxon test (J-L) (*\*p < 0.05, **p < 0.01, ***p < 0.001, ****p < 0.0001.* See also Figure 2.

**Figure S4:**
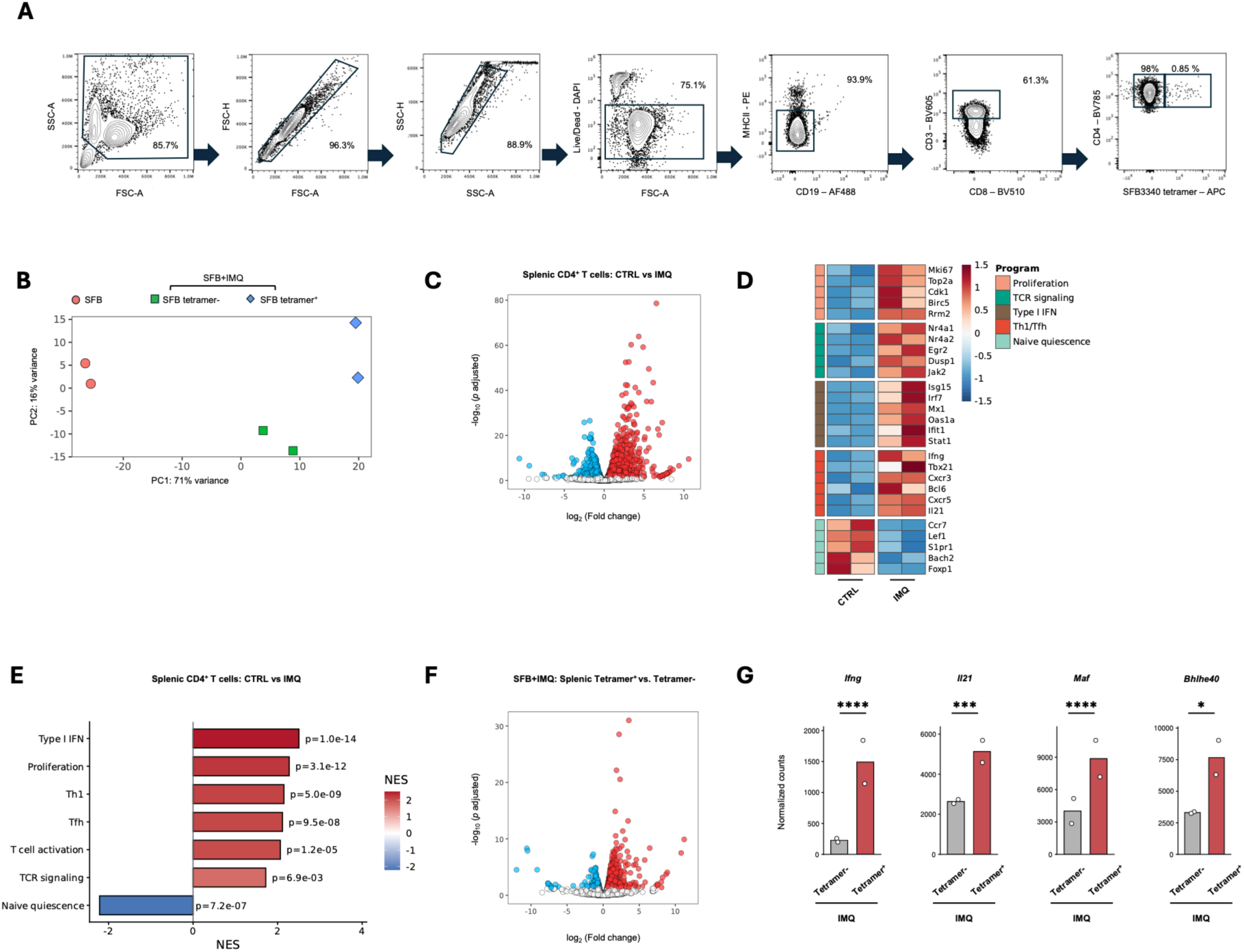
Transcriptional programs of SFB tetramer⁺ CD4⁺ T cells during TLR7-driven autoimmunity. (A) Sorting strategy for SFB tetramer^-^ and SFB tetramer^+^ CD4⁺ T cells from thymus and spleen. For all comparisons in this RNA-sequencing experiment, n = 2 replicates/group, each from 4 pooled mice. (B) PCA plot of total CD4^+^ T cells from SFB-colonized mice and SFB tetramer^-^ and SFB tetramer^+^ cells from SFB-colonized IMQ-treated mice (SFB+IMQ). (C and D) Volcano plot (C) and heatmap (D) of differentially expressed genes (DEGs) in splenic CD4⁺ T cells from healthy control (CTRL) or imiquimod-treated (IMQ) mice. (E) Differentially expressed pathways in healthy CTRL vs. IMQ mice. (F) Volcano plot of DEGs in SFB tetramer^-^ vs. SFB tetramer^+^ cells. (G) Normalized counts of four enriched genes in SFB tetramer^+^ cells. Data shown as means. Statistical significance described in RNA-sequencing analysis methods section. See also Figure 2.

**Figure S5:**
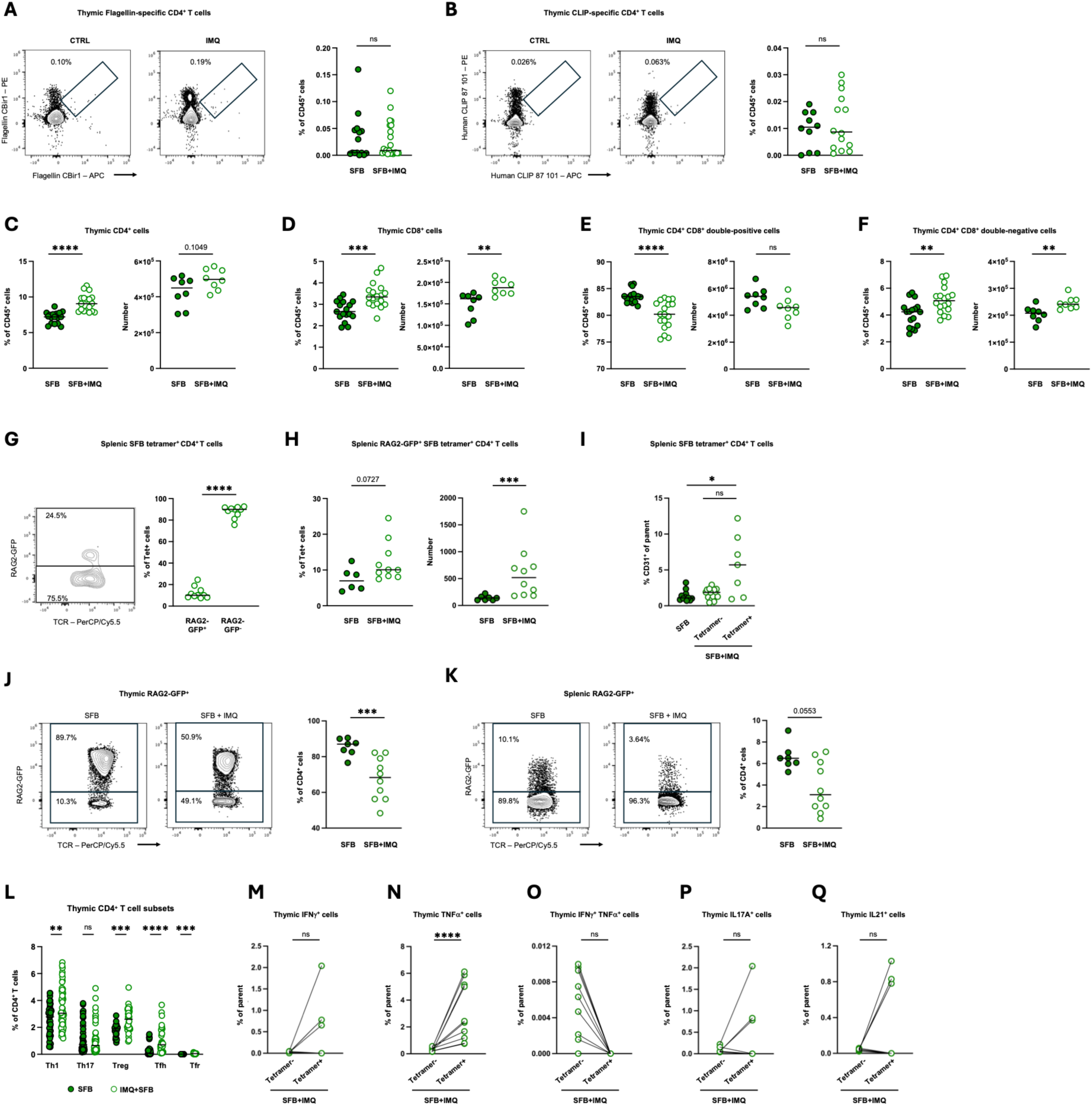
Evidence for *de novo* thymic generation of SFB tetramer⁺ CD4⁺ T cells during TLR7-driven autoimmunity. (A and B) Flagellin tetramer^+^ (A) or CLIP (B) tetramer^+^ CD4⁺ T cells in the thymus of SFB-colonized (SFB) or SFB-colonized imiquimod-treated (SFB+IMQ) mice (n = 10-22). (C-F) Frequencies and counts of thymocyte subsets: CD4 single-positive (C), CD8 single-positive (D), CD4^+^ CD8^+^ double positive (E), and CD4^+^ CD8^+^ double negative (DN) (F) from SFB and SFB+IMQ mice (n = 17-18, n = 8 for all counts). (G) Splenic GFP⁺ cells out of SFB tetramer^+^ CD4⁺ T cells in reporter mice expressing GFP at the *rag2* locus (RAG2-GFP mice) (n = 10). (H) Splenic GFP⁺ SFB tetramer^+^ CD4⁺ T cells in RAG2-GFP mice (n = 6-10). (I) CD31⁺ cells of total splenic CD4⁺ from SFB and SFB tetramer^-^ and SFB tetramer^+^ splenic CD4⁺ T cells from SFB+IMQ mice (n = 7-15). (J-K) GFP⁺ CD4⁺ T cells in thymus (J) and spleen (K) of RAG2-GFP mice (n = 7-10). (L) Effector T cell subsets of total CD4+ T cells in thymus (n = 14-54). (M-Q) IFNγ⁺ (M), TNFα^+^ (N), TNFα^+^ IFNγ⁺ (O), IL-17A⁺ (P), and IL-21^+^ (Q) production in SFB tetramer^-^ and SFB tetramer^+^ CD4+ T cells from thymus of SFB+IMQ mice (n = 10). Data shown as means. Statistical significance was determined by unpaired two-tailed Mann-Whitney U test (A-H, J-L), Kruskal-Wallis test (I), or two-tailed Wilcoxon test (M-Q). *\*p < 0.05, **p < 0.01, ***p < 0.001, ****p < 0.0001.* See also Figure 3.

**Figure S6:**
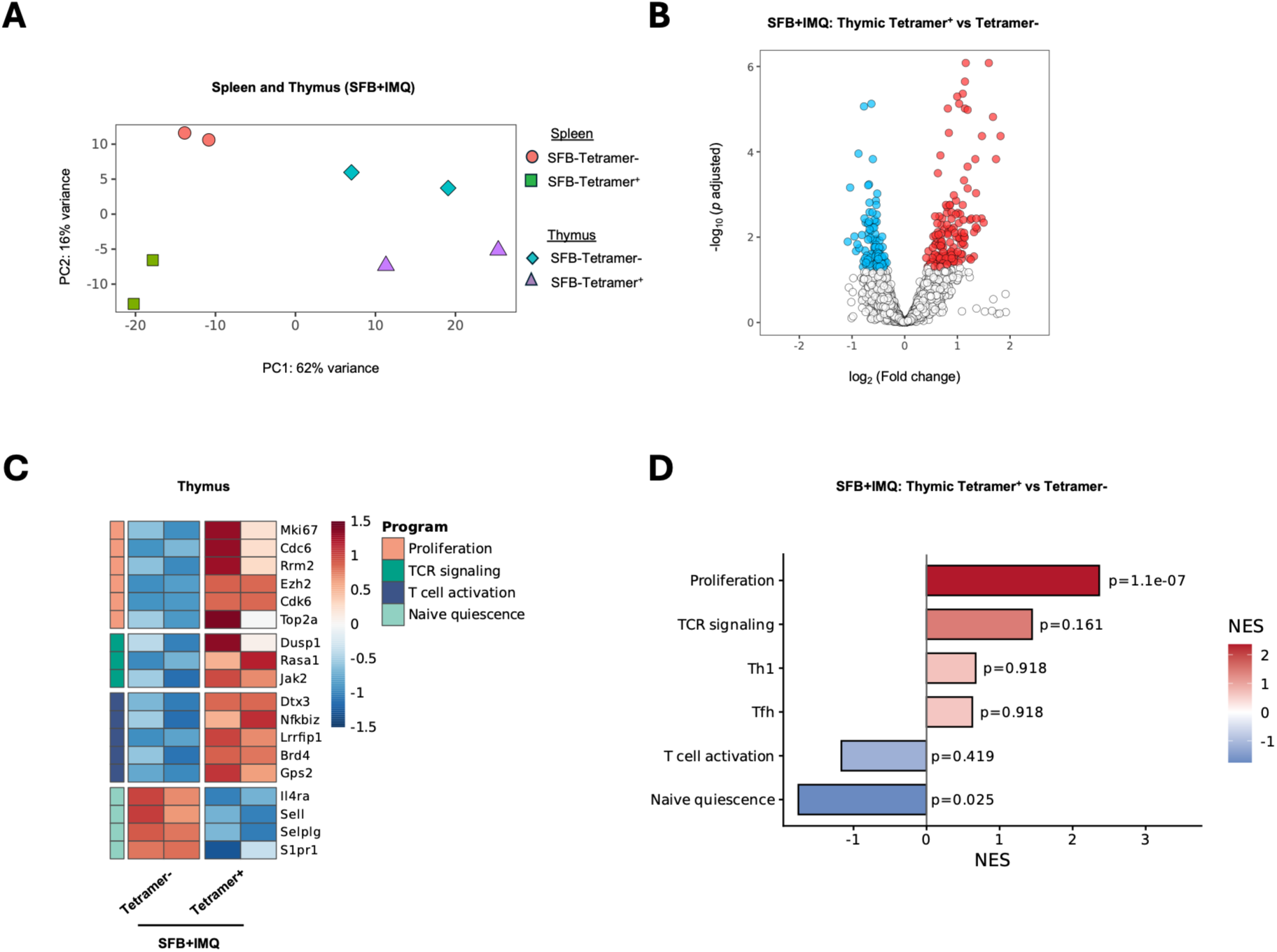
Transcriptional profiling of thymic SFB tetramer⁺ CD4⁺ T cells. (A) PCA of splenic and thymic SFB tetramer^-^ and SFB tetramer^+^ CD4⁺ T cells from SFB-colonized IMQ-treated mice (SFB+IMQ). (B and C) Heatmap (B) and volcano plot (C) of differentially expressed genes between thymic SFB tetramer^-^ and SFB tetramer^+^ CD4+ T cells (n = 2 replicates/group, each from 4 pooled mice). (D) Differentially expressed pathways in SFB tetramer^-^ and SFB tetramer^+^ populations in thymus of SFB+IMQ mice (n = 2 replicates/group, each from 4 pooled mice). See also Figure 3.

**Figure S7:**
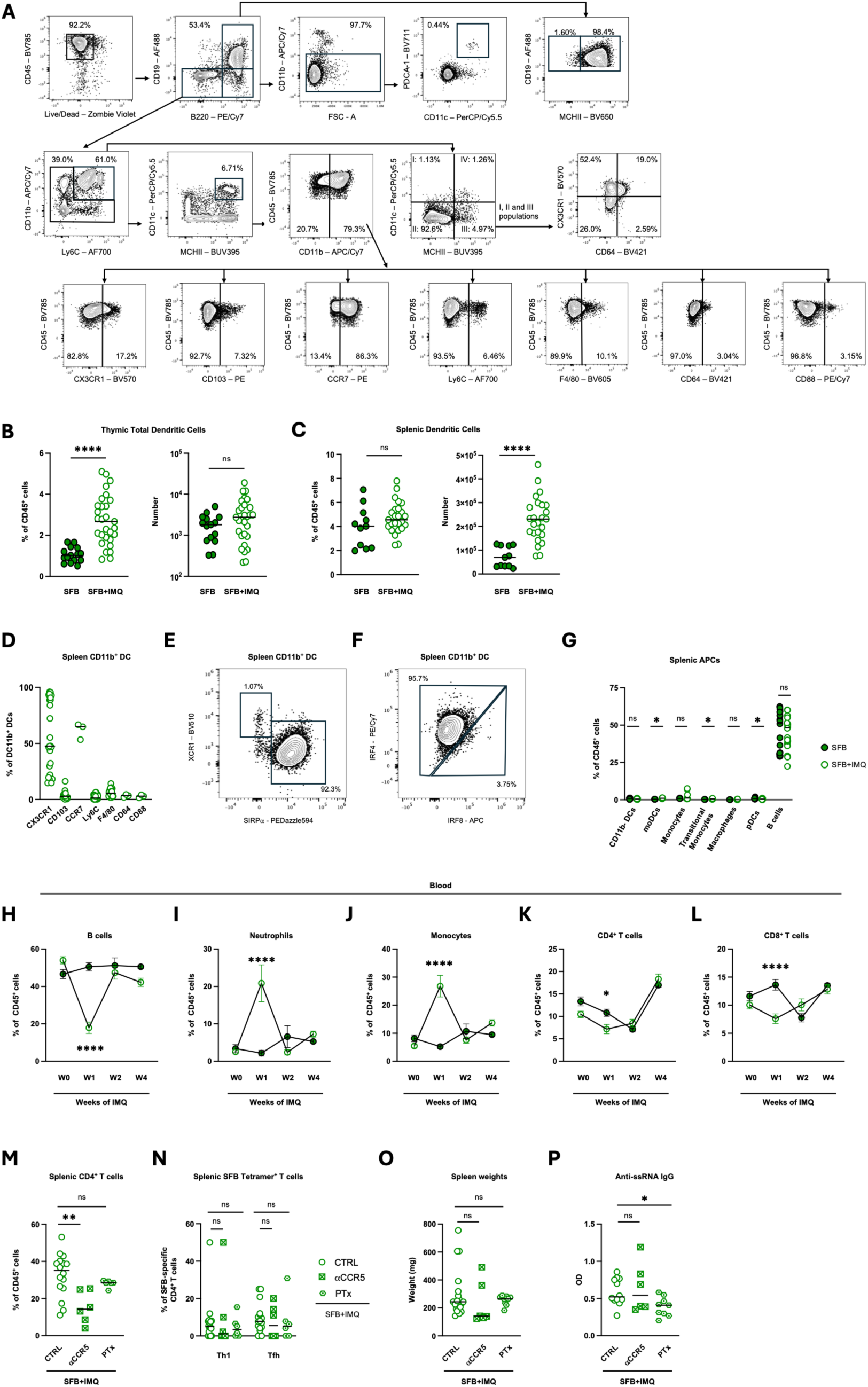
Characterization of antigen-presenting cell populations and intestinal dendritic cell trafficking during TLR7-driven autoimmunity. (A) Gating strategy for antigen-presenting cells and chemokine receptor expression. (B and C) Percentage and counts of total dendritic cells (DCs) in the thymus (B) and spleen of SFB-colonized control (SFB) and IMQ-treated (SFB+IMQ) mice (C) (n = 11-29). (D-F) CD11b+ DC marker gene expression (n = 3-25). (G) Non-DC antigen-presenting cell (APC) frequencies in spleen of SFB and SFB+IMQ mice (n = 4-22). (H-L) Frequencies of B cells (H), neutrophils (I), monocytes (J), CD4+ T cells (K), and CD8+ T cells (L) in blood from weeks 0 to 4 with or without IMQ treatment (n = 15-18). (M-N) Total CD4+ (M) and SFB tetramer^+^ Th1 and Tfh (N) T cell numbers in spleen after anti-CCR5 blockade (aCCR5) or pertussis toxin (PTx) treatment (n = 6-16). (O-P) Spleen weights (O) and serum anti-RNA IgG (P) after aCCR5 or PTx treatment (n = 6-16). Data shown as means. Statistical significance was determined by unpaired two-tailed Mann-Whitney U test (B-C, G), two-way ANOVA (H-L), or Kruskal-Wallis test (M-P). *\*p < 0.05, **p < 0.01, ***p < 0.001, ****p < 0.0001.* See also Figure 5.

**Figure S8:**
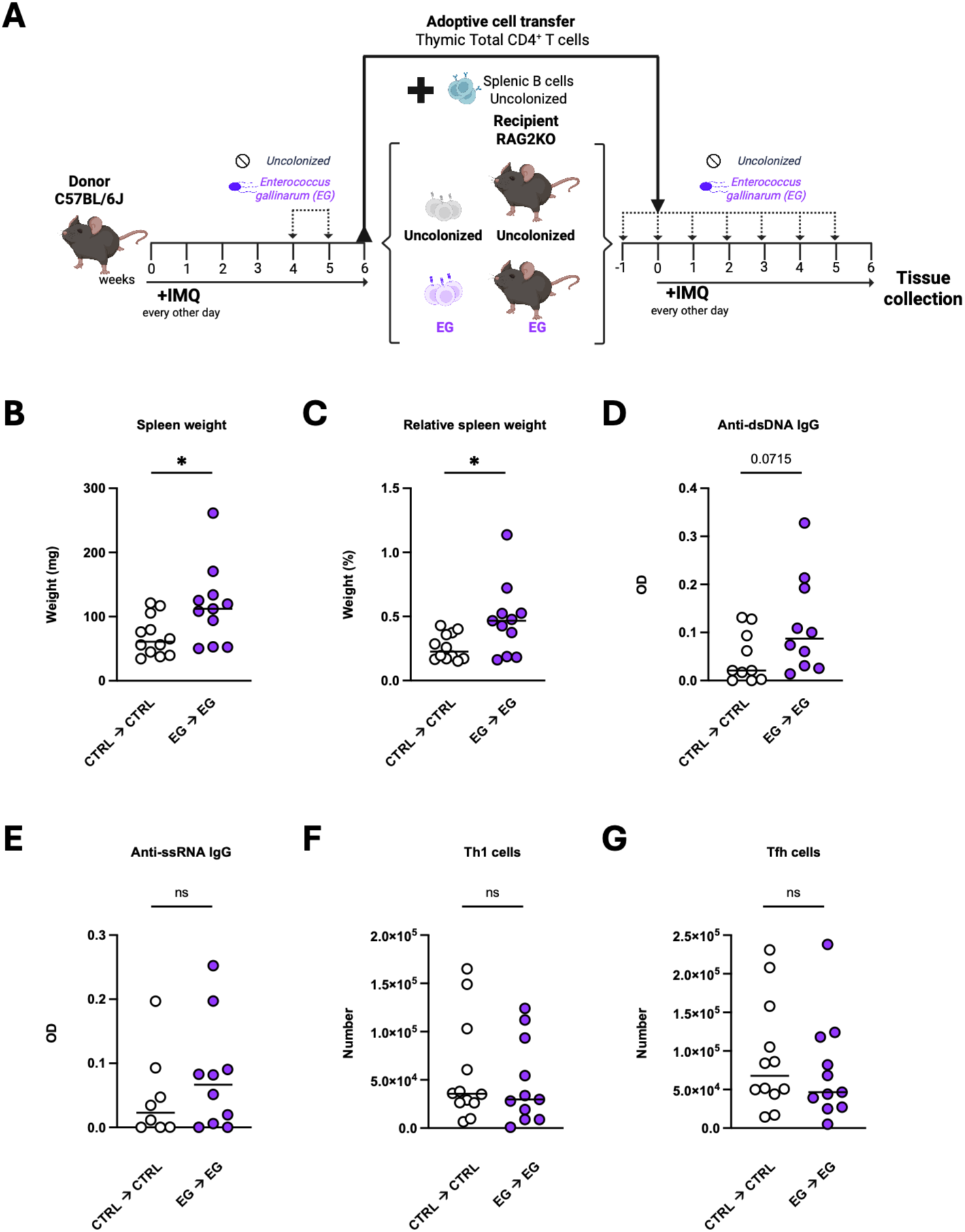
Thymic *E. gallinarum*-reactive CD4⁺ T cells exhibit limited pathogenic potential following adoptive transfer. (A) Experimental schematic of adoptive transfer of thymic CD4⁺ T cells isolated from IMQ-treated wild-type mice either uncolonized (CTRL) or colonized with *Enterococcus gallinarum* (EG). Thymic CD4⁺ T cells were transferred into RAG2KO recipients colonized to match donor microbial status (CTRL → CTRL, EG → EG). Recipient mice were co-transferred with B cells from uncolonized controls and treated with IMQ following adoptive transfer. (B and C) Spleen weight (B) and spleen/body weight ratio (C) in recipients of thymic CD4^+^ T cells (n = 11-12). (D and E) Serum anti-dsDNA IgG (D) and anti-ssRNA IgG (E) (n = 8-10). (F and G) Counts of splenic Th1 (F) and Tfh populations (G) (n = 10-12). Panel A created with BioRender.com. Data shown as means. Statistical significance was determined by unpaired two-tailed Mann-Whitney U test. *\*p < 0.05, **p < 0.01, ***p < 0.001, ****p < 0.0001.* See also Figure 7.

**Figure S9:**
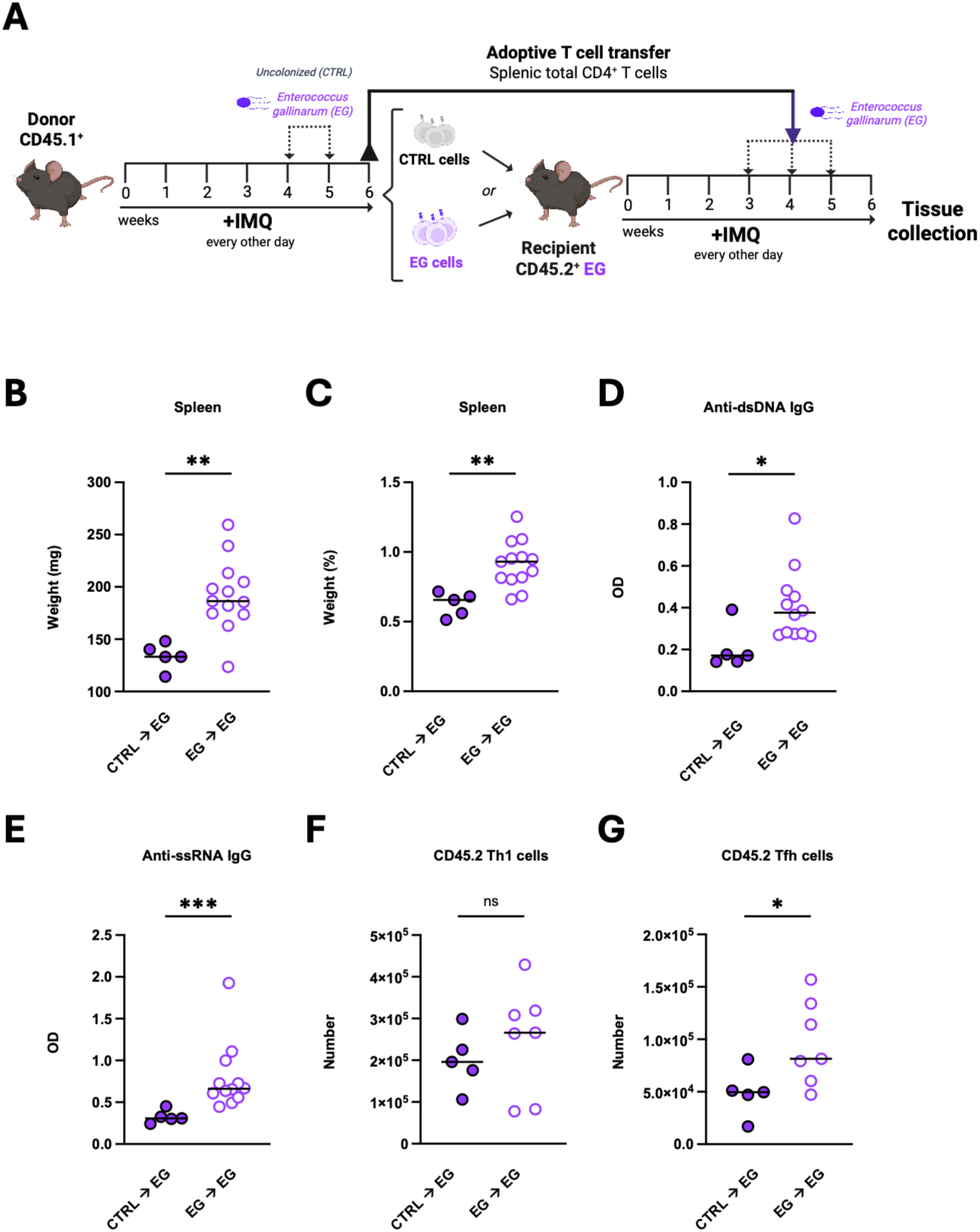
*E. gallinarum*-conditioned CD4^+^ T cells retain pathogenic activity in immunocompetent hosts. (A) Experimental schematic of adoptive transfer of thymic CD4⁺ T cells isolated from IMQ-treated CD45.1⁺ mice either uncolonized (CTRL) or colonized with *Enterococcus gallinarum* (EG). Thymic CD4⁺ T cells were transferred into CD45.2⁺ recipients colonized with *E. gallinarum* (CTRL → EG, EG → EG). Recipient mice were co-transferred with B cells from uncolonized controls and treated with IMQ following adoptive transfer. (B and C) Spleen weight (B) and percentage of total mouse weight (C) in CD45.2⁺ recipients (n = 5-13). (D and E) Serum anti-ssRNA (D) and anti-dsDNA (E) IgG (n= 5-13). (F-G) Recipient Th1 (F) and Tfh (G) cells in spleen (n= 5-7). Panel A was created with Biorender.com. Data shown as means. Statistical significance was determined by unpaired two-tailed Mann-Whitney U test. *\*p < 0.05, **p < 0.01, ***p < 0.001, ****p < 0.0001.* See also Figure 7.

**Figure S10:**
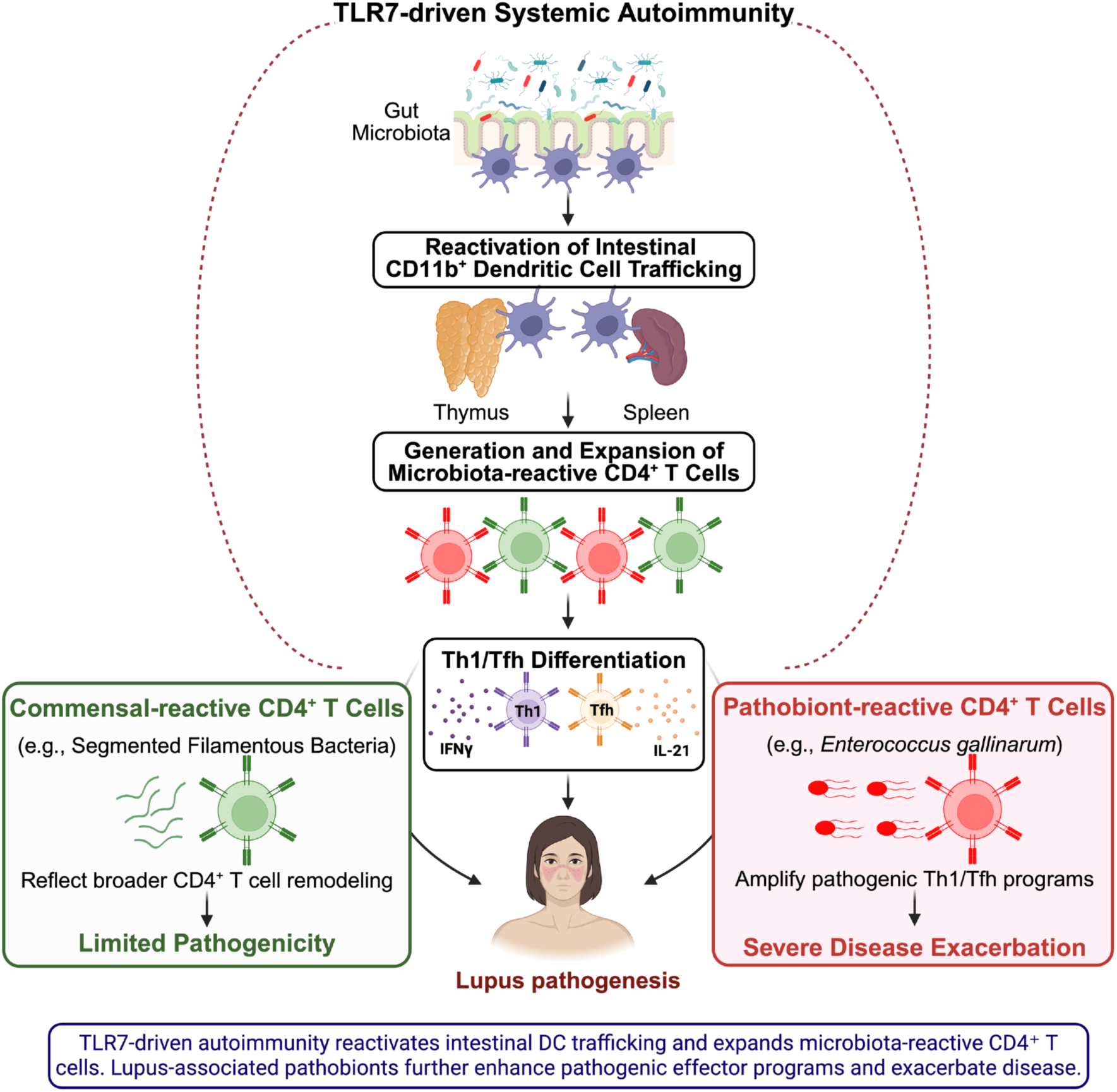
Working model. TLR7-driven autoimmunity reactivates trafficking of CD11b^+^ dendritic cells (DCs) from the gut to systemic organs, including the thymus. CD11b^+^ DC trafficking to the thymus promotes generation and expansion of microbiota-reactive CD4^+^ T cells, which are skewed towards a Tfh/Th1 phenotype. Microbiota-reactive T cells specific to commensals exert limited autoimmune pathogenicity, whereas pathobiont-reactive T cells amplify pathogenic Th1/Tfh programs and exacerbate disease.

## REFERENCES

1. W. E. Ruff, T. M. Greiling, M. A. Kriegel, Host–microbiota interactions in immune-mediated diseases. Nature Reviews Microbiology 2020 18:9 18, 521–538 (2020).

2. M. Kume, J. Din, D. F. Zegarra-Ruiz, Dysregulated Intestinal Host–Microbe Interactions in Systemic Lupus Erythematosus: Insights from Patients and Mouse Models. Microorganisms 2025*, Vol.* 13, *Page 556* **13**, 556 (2025).

3. K. Yao, Y. Xie, J. Wang, Y. Lin, X. Chen, T. Zhou, Gut microbiota: a newly identified environmental factor in systemic lupus erythematosus. Front. Immunol. 14, 1202850 (2023).

4. S. Manfredo Vieira, M. Hiltensperger, V. Kumar, D. Zegarra-Ruiz, C. Dehner, N. Khan, F. R. C. Costa, E. Tiniakou, T. Greiling, W. Ruff, A. Barbieri, C. Kriegel, S. S. Mehta, J. R. Knight, D. Jain, A. L. Goodman, M. A. Kriegel, Translocation of a gut pathobiont drives autoimmunity in mice and humans. Science (1979). 359, 1156–1161 (2018).

5. K. Gronke, M. Nguyen, H. Fuhrmann, N. S. de Souza, J. Schumacher, M. S. Pereira, U. Löschberger, A. Brinkhege, N. J. Becker, Y. Yang, N. Sonnert, S. Leopold, A. L. Martin, L. von Münchow-Klein, C. P. Rodrigues, D. Cansever, R. Hallet, K. Richter, D. A. Schubert, G. M. Daniel, D. Dylus, M. Forkel, D. Schwinge, C. Schramm, S. Redanz, K. G. Lassen, S. M. Vieira, L. Piali, N. W. Palm, C. Bieniossek, M. A. Kriegel, Translocating gut pathobiont Enterococcus gallinarum induces TH17 and IgG3 anti-RNA–directed autoimmunity in mouse and human. Sci. Transl. Med. 17 (2025), doi:10.1126/scitranslmed.adj6294.

6. T. M. Greiling, C. Dehner, X. Chen, K. Hughes, A. J. Iñiguez, M. Boccitto, D. Z. Ruiz, S. C. Renfroe, S. M. Vieira, W. E. Ruff, S. Sim, C. Kriegel, J. Glanternik, X. Chen, M. Girardi, P. Degnan, K. H. Costenbader, A. L. Goodman, S. L. Wolin, M. A. Kriegel, Commensal orthologs of the human autoantigen Ro60 as triggers of autoimmunity in lupus. Sci. Transl. Med. 10 (2018), doi:10.1126/scitranslmed.aan2306.

7. A. Amarnani, C. F. Rivera, M. Cornwell, T. Weinstein, Z. Azad, S. R. S. Gottesman, C. Loomis, A. Lee, N. Ullah, J. Prasad, M. Yi, L. Cooney, B. J. Barnes, N. Gisch, K. V Ruggles, B. Ramkhelawon, G. J. Silverman, A pathogenic gut lipoglycan drives systemic thromboinflammation in lupus nephritis. Ann. Rheum. Dis. (2027), doi:10.1016/j.ard.2026.03.002.

8. D. Azzouz, A. Omarbekova, A. Heguy, D. Schwudke, N. Gisch, B. H. Rovin, R. Caricchio, J. P. Buyon, A. V. Alekseyenko, G. J. Silverman, Lupus nephritis is linked to disease-activity associated expansions and immunity to a gut commensal. Ann. Rheum. Dis. 78, 947–956 (2019).

9. D. F. Zegarra-Ruiz, A. El Beidaq, A. J. Iñiguez, M. Lubrano Di Ricco, S. Manfredo Vieira, W. E. Ruff, D. Mubiru, R. L. Fine, J. Sterpka, T. M. Greiling, C. Dehner, M. A. Kriegel, A Diet-Sensitive Commensal Lactobacillus Strain Mediates TLR7-Dependent Systemic Autoimmunity. Cell Host Microbe 25, 113–127.e6 (2019).

10. S. C. Choi, J. Brown, M. Gong, Y. Ge, M. Zadeh, W. Li, B. P. Croker, G. Michailidis, T. J. Garrett, M. Mohamadzadeh, L. Morel, Gut microbiota dysbiosis and altered tryptophan catabolism contribute to autoimmunity in lupus-susceptible mice. Sci. Transl. Med. 12 (2020), doi:10.1126.

11. R. Duchmann, E. May, M. Heike, P. Knolle, M. Neurath, K. H. Meyer Zum Büschenfelde, T cell specificity and cross reactivity towards enterobacteria,Bacteroides,Bifidobacterium, and antigens from resident intestinal flora in humans. Gut 44, 812–818 (1999).

12. A. N. Hegazy, N. R. West, M. J. T. Stubbington, E. Wendt, K. I. M. Suijker, A. Datsi, S. This, C. Danne, S. Campion, S. H. Duncan, B. M. J. Owens, H. H. Uhlig, A. McMichael, A. Bergthaler, S. A. Teichmann, S. Keshav, F. Powrie, Circulating and Tissue-Resident CD4+ T Cells With Reactivity to Intestinal Microbiota Are Abundant in Healthy Individuals and Function Is Altered During Inflammation. Gastroenterology 153, 1320–1337.e16 (2017).

13. M. Yokogawa, M. Takaishi, K. Nakajima, R. Kamijima, C. Fujimoto, S. Kataoka, Y. Terada, S. Sano, Epicutaneous application of toll-like receptor 7 agonists leads to systemic autoimmunity in wild-type mice: A new model of systemic lupus erythematosus. Arthritis and Rheumatology 66, 694–706 (2014).

14. A. E. Moran, K. L. Holzapfel, Y. Xing, N. R. Cunningham, J. S. Maltzman, J. Punt, K. A. Hogquist, T cell receptor signal strength in Treg and iNKT cell development demonstrated by a novel fluorescent reporter mouse. J. Exp. Med. 208, 1279–89 (2011).

15. Y. Yang, M. B. Torchinsky, M. Gobert, H. Xiong, M. Xu, J. L. Linehan, F. Alonzo, C. Ng, A. Chen, X. Lin, A. Sczesnak, J. J. Liao, V. J. Torres, M. K. Jenkins, J. J. Lafaille, D. R. Littman, Focused specificity of intestinal TH17 cells towards commensal bacterial antigens. Nature 2014 510:7503 510, 152–156 (2014).

16. D. F. Zegarra-Ruiz, D. V. Kim, K. Norwood, M. Kim, W. J. H. Wu, F. B. Saldana-Morales, A. A. Hill, S. Majumdar, S. Orozco, R. Bell, J. L. Round, R. S. Longman, T. Egawa, M. L. Bettini, G. E. Diehl, Thymic development of gut-microbiota-specific T cells. Nature 2021 594:7863 594, 413–417 (2021).

17. J. A. Deane, P. Pisitkun, R. S. Barrett, L. Feigenbaum, T. Town, J. M. M. Ward, R. A. Flavell, S. Bolland, Control of Toll-like Receptor 7 Expression Is Essential to Restrict Autoimmunity and Dendritic Cell Proliferation. Immunity 27, 801–810 (2007).

18. K. A. Hogquist, Y. Xing, F.-C. Hsu, V. S. Shapiro, T Cell Adolescence: Maturation Events Beyond Positive Selection. The Journal of Immunology 195, 1351–1357 (2015).

19. S. J. Szabo, S. T. Kim, G. L. Costa, X. Zhang, C. G. Fathman, L. H. Glimcher, A Novel Transcription Factor, T-bet, Directs Th1 Lineage Commitment. Cell 100, 655–669 (2000).

20. M. Pepper, M. K. Jenkins, Origins of CD4+ effector and central memory T cells. Nature Immunology 2011 12:6 12, 467–471 (2011).

21. A. J. Fike, L. T. Nguyen, O. K. Kumova, A. J. Carey, Characterization of CD31 expression on murine and human neonatal T lymphocytes during development and activation. Pediatric Research 2017 82:1 82, 133–140 (2017).

22. J. Li, Y. Li, J. Y. Yao, R. Jin, M. Z. Zhu, X. P. Qian, J. Zhang, Y. X. Fu, L. Wu, Y. Zhang, W. F. Chen, Developmental pathway of CD4+CD8-medullary thymocytes during mouse ontogeny and its defect in Aire-/-mice. Proc. Natl. Acad. Sci. U. S. A. 104, 18175–18180 (2007).

23. I. Yamashita, T. Nagata, T. Tada, T. Nakayama, CD69 cell surface expression identifies developing thymocytes which audition for T cell antigen receptor-mediated positive selection. Int. Immunol. 5, 1139–1150 (1993).

24. B. D. Coder, H. Wang, L. Ruan, D.-M. Su, Thymic Involution Perturbs Negative Selection Leading to Autoreactive T Cells That Induce Chronic Inflammation. The Journal of Immunology 194, 5825–5837 (2015).

25. R. Thomas, W. Wang, D. M. Su, Contributions of Age-Related Thymic Involution to Immunosenescence and Inflammaging. Immunity & Ageing 2020 17:1 17, 2-(2020).

26. H. J. Wu, I. I. Ivanov, J. Darce, K. Hattori, T. Shima, Y. Umesaki, D. R. Littman, C. Benoist, D. Mathis, Gut-residing segmented filamentous bacteria drive autoimmune arthritis via T helper 17 cells. Immunity 32, 815–827 (2010).

27. M. Guilliams, F. Ginhoux, C. Jakubzick, S. H. Naik, N. Onai, B. U. Schraml, E. Segura, R. Tussiwand, S. Yona, Dendritic cells, monocytes and macrophages: A unified nomenclature based on ontogeny. Nat. Rev. Immunol. 14, 571–578 (2014).

28. S. Nowotschin, A. K. Hadjantonakis, Use of KikGR a photoconvertible green-to-red fluorescent protein for cell labeling and lineage analysis in ES cells and mouse embryos. BMC Developmental Biology 2009 9:1 9, 49-(2009).

29. A. M. Morton, E. Sefik, R. Upadhyay, R. Weissleder, C. Benoist, D. Mathis, Endoscopic photoconversion reveals unexpectedly broad leukocyte trafficking to and from the gut. Proc. Natl. Acad. Sci. U. S. A. 111, 6696–701 (2014).

30. S. Galván-Peña, Y. Zhu, B. S. Hanna, D. Mathis, C. Benoist, A dynamic atlas of immunocyte migration from the gut. Sci. Immunol. 9, 672 (2024).

31. J. Ni, Y.-N. Zhu, X.-G. Zhong, Y. Ding, L.-F. Hou, X.-K. Tong, W. Tang, S. Ono, Y.-F. Yang, J.-P. Zuo, The chemokine receptor antagonist, TAK-779, decreased experimental autoimmune encephalomyelitis by reducing inflammatory cell migration into the central nervous system, without affecting T cell function. Br. J. Pharmacol. 158, 2046–56 (2009).

32. M. A. Linterman, R. J. Rigby, R. K. Wong, D. Yu, R. Brink, J. L. Cannons, P. L. Schwartzberg, M. C. Cook, G. D. Walters, C. G. Vinuesa, Follicular helper T cells are required for systemic autoimmunity. Journal of Experimental Medicine 206, 561–576 (2009).

33. S. Crotty, T Follicular Helper Cell Differentiation, Function, and Roles in Disease. Immunity 41, 529–542 (2014).

34. I. I. Ivanov, K. Atarashi, N. Manel, E. L. Brodie, T. Shima, U. Karaoz, D. Wei, K. C. Goldfarb, C. A. Santee, S. V. Lynch, T. Tanoue, A. Imaoka, K. Itoh, K. Takeda, Y. Umesaki, K. Honda, D. R. Littman, Induction of Intestinal Th17 Cells by Segmented Filamentous Bacteria. Cell 139, 485–498 (2009).

35. Y. Goto, C. Panea, G. Nakato, A. Cebula, C. Lee, M. G. Diez, T. M. Laufer, L. Ignatowicz, I. I. Ivanov, Segmented filamentous bacteria antigens presented by intestinal dendritic cells drive mucosal Th17 cell differentiation. Immunity 40, 594–607 (2014).

36. F. E. Mercier, D. B. Sykes, D. T. Scadden, Single Targeted Exon Mutation Creates a True Congenic Mouse for Competitive Hematopoietic Stem Cell Transplantation: The C57BL/6-CD45.1STEM Mouse. Stem Cell Reports 6, 985–992 (2016).

37. R. L. Reinhardt, H. E. Liang, R. M. Locksley, Cytokine-secreting follicular T cells shape the antibody repertoire. Nature Immunology 2009 10:4 10, 385–393 (2009).

38. Z. Hao, K. Rajewsky, Homeostasis of Peripheral B Cells in the Absence of B Cell Influx from the Bone Marrow. Journal of Experimental Medicine 194, 1151–1164 (2001).

39. H. Nagaoka, G. Gonzalez-Aseguinolaza, M. Tsuji, M. C. Nussenzweig, Immunization and Infection Change the Number of Recombination Activating Gene (Rag)-Expressing B Cells in the Periphery by Altering Immature Lymphocyte Production. Journal of Experimental Medicine 191, 2113–2120 (2000).

40. B. Bushnell, BBMap: A Fast, Accurate, Splice-Aware Aligner https://sourceforge.net/projects/bbmap/ (2014) (available at https://sourceforge.net/projects/bbmap/).

41. R. Patro, G. Duggal, M. I. Love, R. A. Irizarry, C. Kingsford, Salmon provides fast and bias-aware quantification of transcript expression. Nature Methods 2017 14:4 14, 417–419 (2017).

42. A. Frankish, M. Diekhans, A. M. Ferreira, R. Johnson, I. Jungreis, J. Loveland, J. M. Mudge, C. Sisu, J. Wright, J. Armstrong, I. Barnes, A. Berry, A. Bignell, S. Carbonell Sala, J. Chrast, F. Cunningham, T. Di Domenico, S. Donaldson, I. T. Fiddes, C. García Girón, J. M. Gonzalez, T. Grego, M. Hardy, T. Hourlier, T. Hunt, O. G. Izuogu, J. Lagarde, F. J. Martin, L. Martínez, S. Mohanan, P. Muir, F. C. P. Navarro, A. Parker, B. Pei, F. Pozo, M. Ruffier, B. M. Schmitt, E. Stapleton, M. M. Suner, I. Sycheva, B. Uszczynska-Ratajczak, J. Xu, A. Yates, D. Zerbino, Y. Zhang, B. Aken, J. S. Choudhary, M. Gerstein, R. Guigó, T. J. P. Hubbard, M. Kellis, B. Paten, A. Reymond, M. L. Tress, P. Flicek, GENCODE reference annotation for the human and mouse genomes. Nucleic Acids Res. 47, D766– D773 (2019).

43. C. Soneson, M. I. Love, M. D. Robinson, Differential analyses for RNA-seq: transcript-level estimates improve gene-level inferences. F1000Research 2016 4:1521 **4**, 1521 (2016).

44. M. I. Love, W. Huber, S. Anders, Moderated estimation of fold change and dispersion for RNA-seq data with DESeq2. Genome Biology 2014 15:12 15, 550-(2014).

45. O. T. Chan, L. G. Hannum, A. M. Haberman, M. P. Madaio, M. J. Shlomchik, A novel mouse with B cells but lacking serum antibody reveals an antibody-independent role for B cells in murine lupus. J. Exp. Med. 189, 1639–48 (1999).

